# A 3D microtumour system that faithfully represents ovarian cancer minimal residual disease

**DOI:** 10.1101/2023.07.15.549155

**Authors:** Xingyun Yang, Mara Artibani, Yongcheng Jin, Aneesh Aggarwal, Yujia Zhang, Sandra Muñoz-Galvan, Ellina Mikhailova, Lena Rai, Nobina Mukherjee, Ravinash Krishna Kumar, Ashwag Albukhari, Linna Zhou, Ahmed Ashour Ahmed, Hagan Bayley

## Abstract

**Background:** Bulk cancer and minimal residual disease (MRD) are characterised by different molecular drivers and therefore necessitate different therapeutic strategies. However, there are currently no 3D models that can faithfully recapitulate MRD *ex vivo* for therapy development.

**Methods:** A microfluidic technique was implemented to construct 3D microtumours, in which tumour cells, either by themselves or with fibroblasts, were encapsulated in viscous hydrogels. The 3D microtumours were analysed for their response to first-line chemotherapeutics and characterised through RNA-Seq, by comparing them to both 2D cultures and clinical samples.

**Results:** Our microfluidic platform guarantees the fabrication of 3D microtumours of tailorable size and cell content, which recreate key features of tumours such as hypoxia, characteristic organization of the cytoskeleton and a dose-response to chemotherapeutics close to the physiological range. The 3D microtumours were also used to examine non-genetic heterogeneity in ovarian cancer and could fully reflect the recently described “Oxford Classic” five molecular signatures.

The gene expression profile of 3D microtumours following chemotherapy treatment closely resembled that of MRD in ovarian cancer patients, showing the upregulation of genes involved in fatty acid metabolism. We demonstrate that these 3D microtumours are ideal for drug development by showing how they support the identification of a promising inhibitor of fatty acid oxidation, perhexiline, which specifically targets chemotherapy-resistant MRD ovarian cancer cells and not bulk cancer cells.

**Conclusion:** We have obtained the first 3D model of ovarian cancer MRD by using microtumours generated through microfluidics. This system is ideal for high-throughput drug screening and, given its versatility, it can be readily extended to additional types of cancer, as well as accommodate multiple cell types to generate complex tumour microenvironments.

## Background

Drug resistance is responsible for up to 90% of cancer-related deaths and can operate through several different mechanisms^1^. A particularly challenging group of treatment-resistant cells is represented by Minimal Residual Disease (MRD), the microscopic clusters of malignant cells that remain in patients after complete clinical/radiological response and are capable of reinitiating tumours^2^. Investing in therapeutics that specifically target MRD could help delay or prevent relapses altogether, moving us a step closer to the chronic management or potential eradication of cancer^3^.

However, this long-sought goal has been difficult to achieve, especially in solid tumours: isolating and characterising these cells directly from patients can be quite challenging. Because of this, we lack deep knowledge of MRD biology as well as appropriate experimental models that could be used as screening platforms for effective compounds.

One of the few solid tumours where direct characterisation of clinical MRD has been achieved is ovarian cancer^4^. After obtaining the first in-depth transcriptomic characterisation of MRD from patients, we aimed to develop a faithful and physiologically-relevant model of MRD that could be used for drug screening. We considered mouse, 2D and 3D models, all of which have different balances of feasibility, accuracy and cost.

Although mouse models exist to study metastatic ovarian cancer, their reproductive physiology^5^ as well as omental anatomy (one of the most common MRD sites)^6^ differ from humans and, further, this species does not develop spontaneous ovarian tumours. Conventional 2D cell culture has been a standard *in vitro* model for decades. However, under these conditions, cell morphologies and cell bioactivities deviate from those found *in vivo*^7^. For instance, cells in 2D culture lose diverse phenotypes, have altered cell signalling, rely on different metabolic pathways, and have an unlimited supply of nutrients and oxygen^8^, which means they may not provide biologically meaningful responses to chemotherapeutics. Moreover, a transcriptomic study of single cell-derived spheroids from ovarian cancer ascites has shown that although 2D monolayers support proliferation and tumour growth cascades, 3D spheroids additionally capture aspects of cholesterol and lipid metabolism, which are features of metastatic disease^9^. These pathways are implicated in the lipid signature we observed in ovarian cancer MRD^4^; thus, in this context, 3D models are essential for drug discovery.

Another aspect to be considered is that cancers typically organise in 3D, creating a hypoxic setting with intimate intercellular signalling, and ultimately achieve anchorage-independent growth. 3D cancer cultures offer extensive cell-cell and cell-ECM (extracellular matrix) interactions^10^, and are understood to preserve the cell polarity, morphology, gene expression and topology seen *in vivo*^8, 11, 12^. They also offer the prospect to co-culture vascular and stromal elements and investigate heterotypic interactions^13^. This is a crucial prerequisite for our model, since environment-mediated drug resistance has been shown to be a major contributor to MRD^14^.

3D cell culture technologies can be classified as scaffold-free or scaffold-based. Scaffold-free systems include multi-cellular aggregates formed using the hanging-drop method^15, 16^, suspension plates^17, 18^, silicone micro-moulds^19^ (which can prevent cell adhesion), or spinner flasks^20^. However, the formation of cell aggregates with a large number of cells (>500 µm) requires prolonged incubation over several days. Additionally, 3D microtumours fabricated by these methods often exhibit significant variation in size, limiting their application in the screening of therapeutics.

Scaffold-based systems employ biocompatible materials, such as hydrogels, as structural supports for cell culture^8, 13^. Cells proliferate in the scaffolds and establish cell-cell and cell-ECM interactions, displaying natural 3D structures instead of flattening out as they do in 2D culture^21^. Despite all these advantages, the application of 3D cell culture models to drug screening has been constrained by long fabrication times, poor repeatability and low productivity^8^.

Microfluidics is a promising tool for dealing with various unmet needs in 3D cell culture. Based on the immiscibility of aqueous and oil phases, discrete aqueous droplets of uniform size and composition can be generated by microfluidics, in which cells can be encapsulated in a highly reproducible and high-throughput manner for subsequent 3D culture. For example, microfluidics has been adopted to create organoids or tumour spheroids for predicting drug responses^22^, investigating tumour vascularization^23^, and for studying hair follicle regeneration with stem cell patterning^24^. In the present work, a microfluidics platform was established and optimised to fabricate scaffold-based 3D microtumours, with readily customisable size, morphology and hydrogel choice. 3D microtumours of various size were produced by this approach, and key characteristics, such as cell viability and hypoxic core formation, were determined. First-line chemotherapeutics screening was performed on the 3D microtumours to demonstrate their utility in pharmacology. Finally, 3D microtumours were successfully used as a drug screening platform for ovarian cancer MRD. The structures showed the same molecular signatures observed in clinical MRD and led not only to the identification of a very promising therapeutic agent but also to significant insight into resistance mechanisms.

## Results

### Microfluidic platform generates 3D microtumours of tailorable size, cell content, and shape

The 3D microtumours, tumour cells encapsulated in biocompatible hydrogels, were fabricated by using a surfactant-free, droplet-based microfluidic platform (Figure 1A). Specifically, ovarian cancer cells OVCAR-5/RFP were mixed with Matrigel to form the bioink (~3.5×10^7^ cells mL^−1^). The bioink and tetradecane oil were loaded into two separate syringes and pumped into a 3-channel polydimethylsiloxane (PDMS) microfluidic chip (Figure S1A), in which the two immiscible liquids met and produced 3D microtumours separated by oil. The microfluidic fabrication was performed at 8°C to prevent the Matrigel gelation or viscosity change. At least one hundred 3D microtumours were produced within 3 minutes and temporarily stored in a polytetrafluoroethylene (PTFE) exit tube (inner diameter = 900 µm) (Figure S1B) connected to the microfluidic chip. The exit tube was then incubated at 37°C for 2 h to complete gelation of the Matrigel. The temperature and time required for gelation varies with the hydrogel, which included collagen, agarose and silk fibroin (Table S1, Figure S1C, see Methods for more details). In addition to OVCAR-5/RFP, we have also successfully fabricated 3D microtumours with other tumour cell lines (OVCAR5, OVCAR8, Kuramochi, MDA-MB-231, HeLa) and similar constructs from normal tissue cell lines (fibroblasts, adipocytes, embryonic kidney cells) (Table S2). Cells proliferated within the 3D microtumours, which increased their density by D2 (Figure 1A). Long term culture of 3D microtumours for up to 9 weeks has been achieved and no major morphological changes have been observed (Figure S1D).

**Figure 1.**
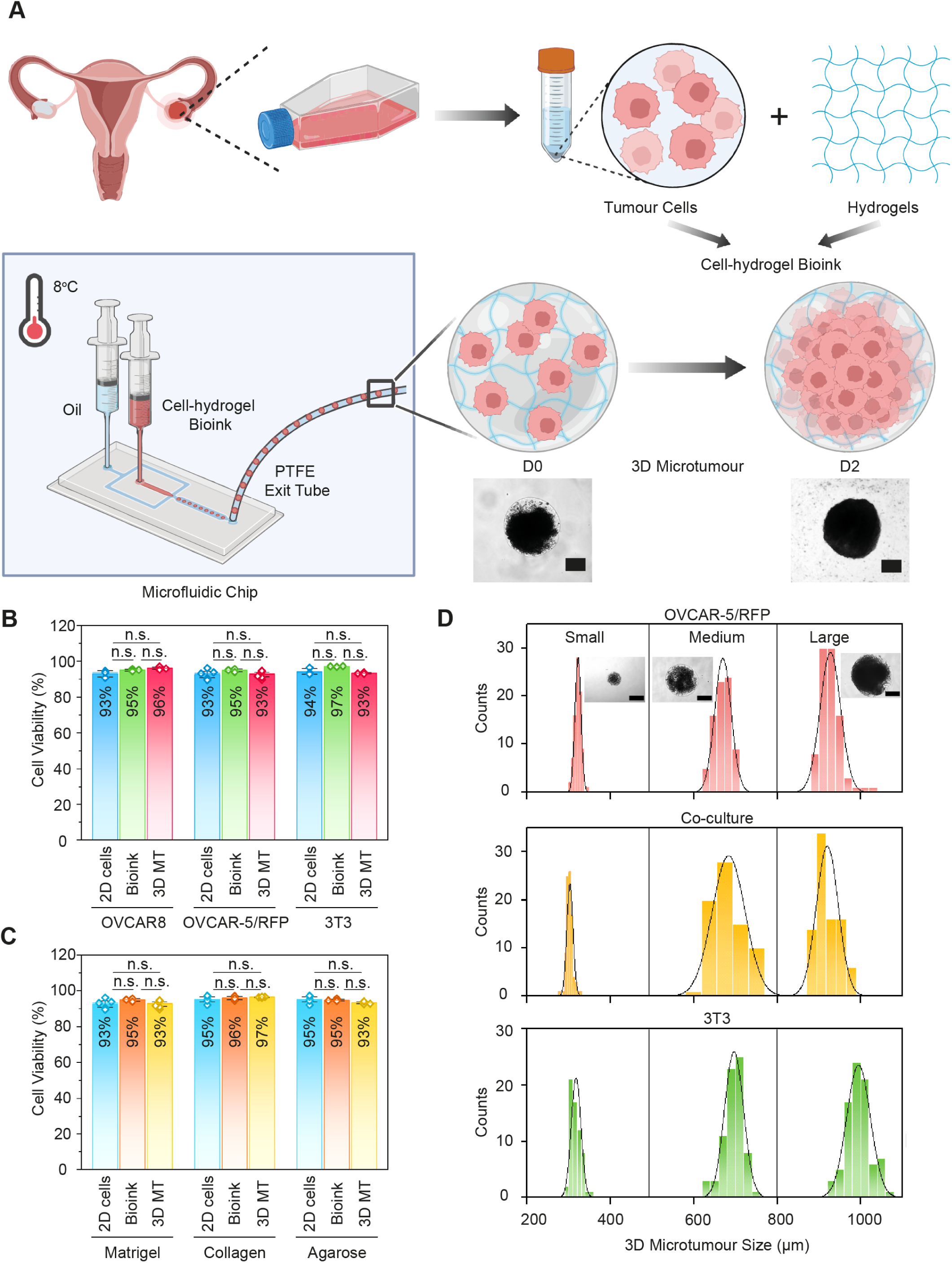
Schematic illustration and features of 3D microtumour fabrication by the microfluidic platform. A) 2D monolayer cells (ovarian cancer cells) were harvested and mixed with hydrogel (Matrigel) to form a bioink. The cell-hydrogel bioink and tetradecane oil were pumped into a PDMS microfluidic chip at 8°C by syringe pumps. At least one hundred 3D microtumours separated by oil were generated within 3 minutes and temporarily stored in a PTFE exit tube. Upon complete gelation, the 3D microtumours were transferred from the PTFE tube to a medium-containing culture plate for further use. Inset: Microscope images of 3D microtumours composed of OVCAR-5/RFP and Matrigel on D0 and D2 of fabrication. Scale bar = 300 µm. The diagrams were created with BioRender.com. B) Cell viability at three stages of microfluidic fabrication: 2D cells harvested from flasks, bioink made from cells and Matrigel, and 3D microtumours after microfluidic fabrication (3D MT). Data for three cell lines are shown: OVCAR8, OVCAR-5/RFP and 3T3 fibroblasts (n = 3 to 6). C) Cell viability at the three stages of microfluidic fabrication for OVCAR8 cells with three hydrogels: Matrigel, collagen and agarose (n = 3 to 6). D) Size distributions of Matrigel 3D microtumours composed of OVCAR-5/RFP tumour cells, a 50: 50 mixture of OVCAR-5/RFP cells and 3T3 cells (co-culture) and 3T3 fibroblast cells (n = 62 to 90). Inset: microscope images of 3D microtumours generated with PTFE exit tubes of ID = 300, 650 and 900 µm. Scale bar = 300 µm.

To examine if the microfluidic fabrication process had caused damage to the cells, viability assays were performed at three stages of microfluidic fabrication: 2D cells harvested from culture flasks, cell-hydrogel bioink, and 3D microtumours after microfluidic fabrication. Cell viabilities of >90% were achieved with no significant differences observed among all the three stages, proving that the microfluidic fabrication does not harm the cells. High cell viability was maintained with a variety of cells (Figure 1B) and different hydrogels (Figure 1C, Table S3).

The microfluidic platform offers flexibility in adjusting the size, cell content and shape of 3D microtumours, which is hard to achieve with other fabrication methods. By simply using PTFE exit tubes with different inner diameter, 3D microtumours of different sizes (small, medium, and large) can be prepared. Cell contents were managed by adjusting the ratio of different cell types during bioink preparation. Three types of microtumours were fabricated: 1) OVCAR-5/RFP tumour cells only; 2) 50: 50 mixture of OVCAR-5/RFP tumour cells and 3T3 fibroblast cells (co-culture); and 3) 3T3 fibroblast cells only. For each, the sizes of >60 microtumours were determined on the day of fabrication. A histogram showed a narrow size distribution of 3D microtumours from the same batch with 2-6% deviation (Figure 1D, Table S4). The narrow size distribution is essential for reliable and reproducible drug screening experiments, since the cell population within each microtumour depends on its size. Furthermore, different 3D microtumour shapes (sphere, ellipsoid, and rod, Figure S1E) were created by varying the flowrate ratio of bioink and oil.

### 3D microtumours recapitulate key physiological features of tumours

Certain physical and biochemical characteristics of tumour cells are particularly difficult to recreate *in vitro*, which leads to the use of sub-optimal models for drug screening and eventually disappointing results from clinical trials^6^. Among these critical, and hard to model, features of tumours we find low oxygen tension, or hypoxia, and cytoskeleton organisation.

Hypoxia contributes to reshaping of the tumour microenvironment and the development of immunosuppression and chemoresistance^25^, however, 2D monolayer cells lack the gradients of oxygen required to produce hypoxia. Hypoxic cores are commonly observed in tumours with diameters larger than 400-500 µm due to the deficient oxygen and nutrient levels towards the centres of tumours^26, 27^. Nevertheless, the initial size of microtumours prepared by prevailing methods usually falls within the range of 100-300 µm^20, 28^. With the microfluidic platform developed in this work, 3D microtumours with larger initial sizes can conveniently be produced. Significantly, hypoxic cores were observed in large 3D microtumours (size = 825 µm) just 1 day after fabrication (Figure 2A), while they were absent in small and medium size microtumours (size = 274 and 424 µm), which is consistent with the published literature^26, 27^. We also confirmed the expression of key hypoxia genes through immunofluorescence (Figure 2B, C) in older microtumours: at day 10 after fabrication, the hypoxic core had expanded, with expression of P4HA1, VEGFA and NDRG1 observed very strongly in the middle of the structure and, more faintly, in some cells closer to the edges. 3D spheroids prepared with other methods, such as the forced aggregation method, took 11-21 days to grow to a comparable size (>800 µm) and generate hypoxic cores^27, 29^. Therefore, hypoxic characteristics can be recapitulated in one-tenth of the time with the microfluidic platform.

**Figure 2.**
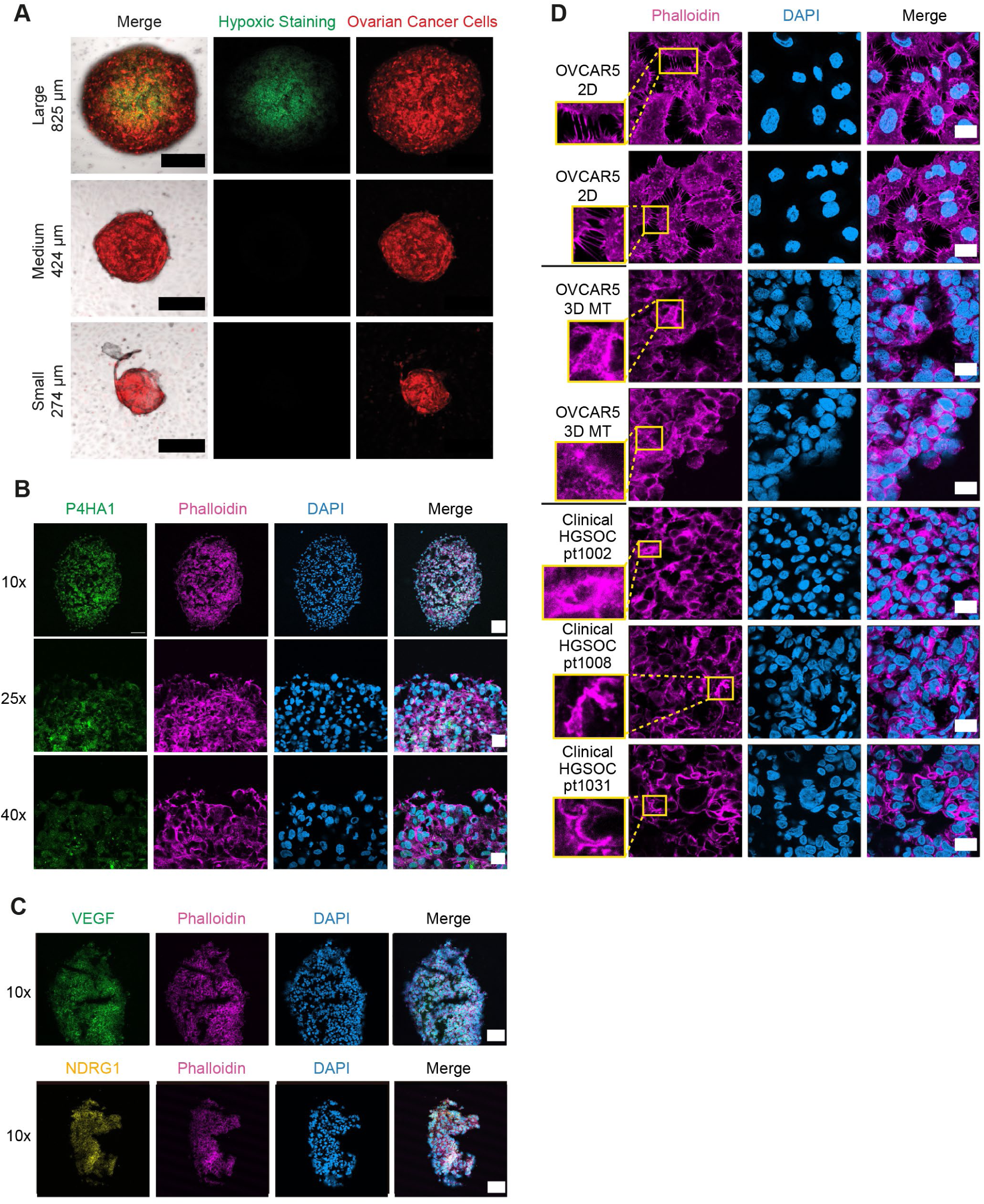
3D microtumours recapitulate key physiological features of cancers. A) Epifluorescence images of hypoxia staining (green) for large, medium and small 3D microtumours composed of OVCAR-5/RFP ovarian cancer cells (red) and Matrigel. Scale bar = 300 µm. B) Confocal images of 3D microtumours composed of OVCAR5 ovarian cancer cells and Matrigel at D10 stained with P4HA1 (green), phalloidin (pink), DAPI (blue). Scale bar = 100 µm for 10x, 30 µm for 25x, and 20 µm for 40x magnification. C) Confocal images of 3D microtumours composed of OVCAR5 ovarian cancer cells and Matrigel at D10 stained with VEGF (green), NDRG1 (yellow), phalloidin (pink), DAPI (blue). Scale bar = 100 µm. D) Confocal images of 2D cultures (OVCAR5), 3D microtumours (3D MT, OVCAR5) at D10, and clinical samples stained with phalloidin (pink), DAPI (blue). Scale bar = 20 µm. Inset: zoom-in images for regions of interest stained with phalloidin.

Another key feature of tumour cells is their cytoskeleton organisation, which plays a crucial role in cell motility, and therefore, in invasion and metastasis. More specifically, actin filaments can act at different levels, from providing a connection with the ECM, to being mechanosensors and signalling scaffolds^30^, all of which are altered in cancer.

Several studies have reported that the actin patterns and dynamics observed in living tumours are not recreated in 2D cultures or most 3D systems either^31^, especially when it comes to stress-fibre structures^32^. To investigate whether our microtumours could recapitulate the same actin distribution observed in ovarian cancer clinical samples, we used phalloidin staining on OVCAR5 cells grown in 2D cultures, OVCAR5 3D microtumours and High Grade Serous Ovarian Cancer (HGSOC) pre-chemotherapy biopsies (Figure 2D). While the 2D cells are characterised by many thin filipodia, a rich network of thick stress fibre-like actin bundles is present in both 3D microtumours and HGSOC samples (which were also stained for E-cadherin to rule out any stromal contamination, Figure S2).

To complement these imaging experiments with a full transcriptomics analysis, we conducted RNA-Seq on both 2D cultures and 3D microtumours produced from three ovarian cancer cell lines (OVCAR5, OVCAR8, OVCAR-5/RFP). Regardless of the cell line used, the microtumours overexpressed genes that fall into several biological processes related to hypoxia (such as a 16-fold enrichment for the positive regulation of VEGF production) and cell motility (Figure 3A). We then analysed the expression of a nine-gene ovarian cancer-specific hypoxic signature^33^ across all our samples: for OVCAR5 and OVCAR-5/RFP almost all genes were consistently upregulated in the microtumours (Figure 3B, C), while for OVCAR8 the upregulation was restricted to VEGF and NDRG1 (Figure S3A, B).

**Figure 3.**
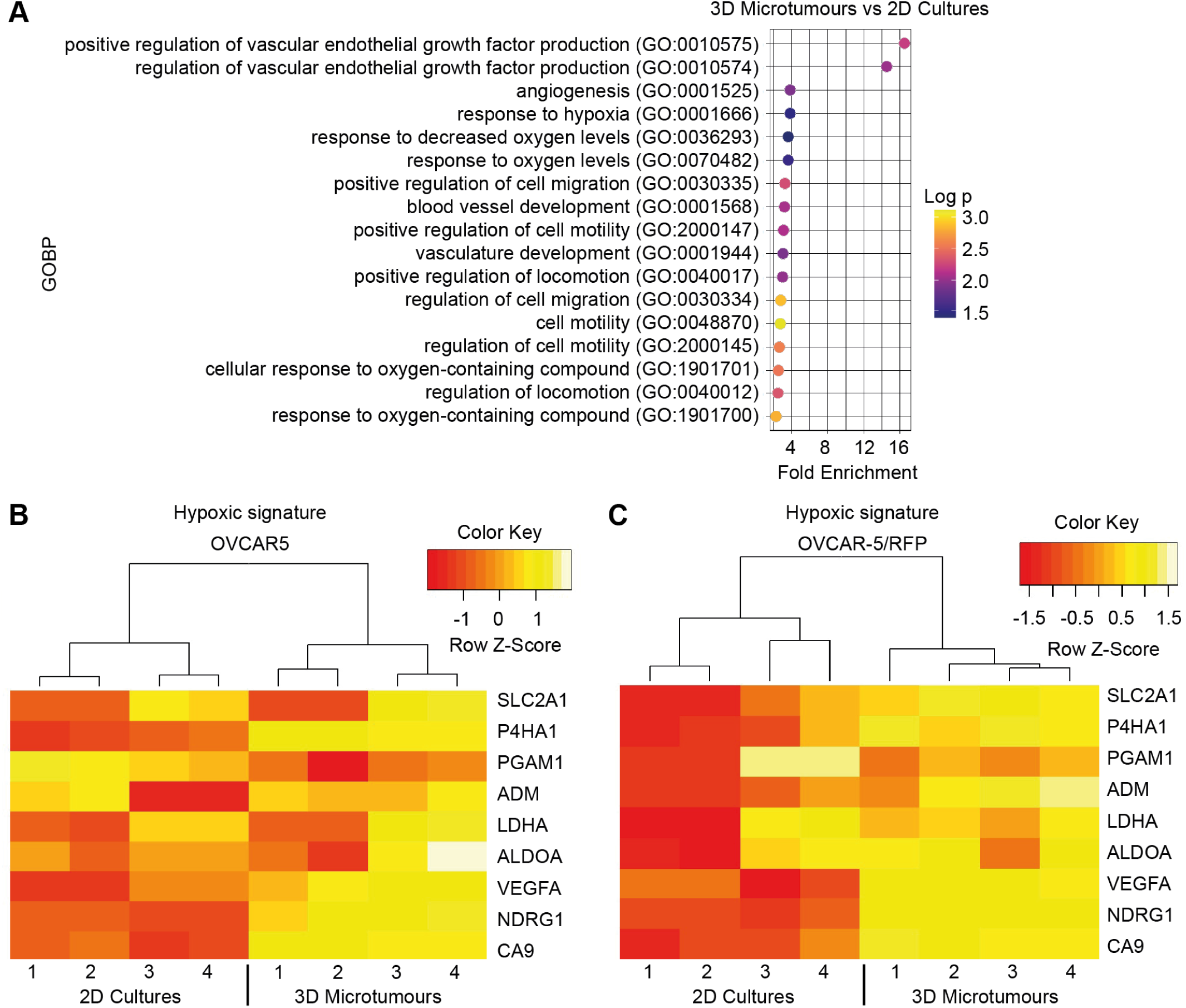
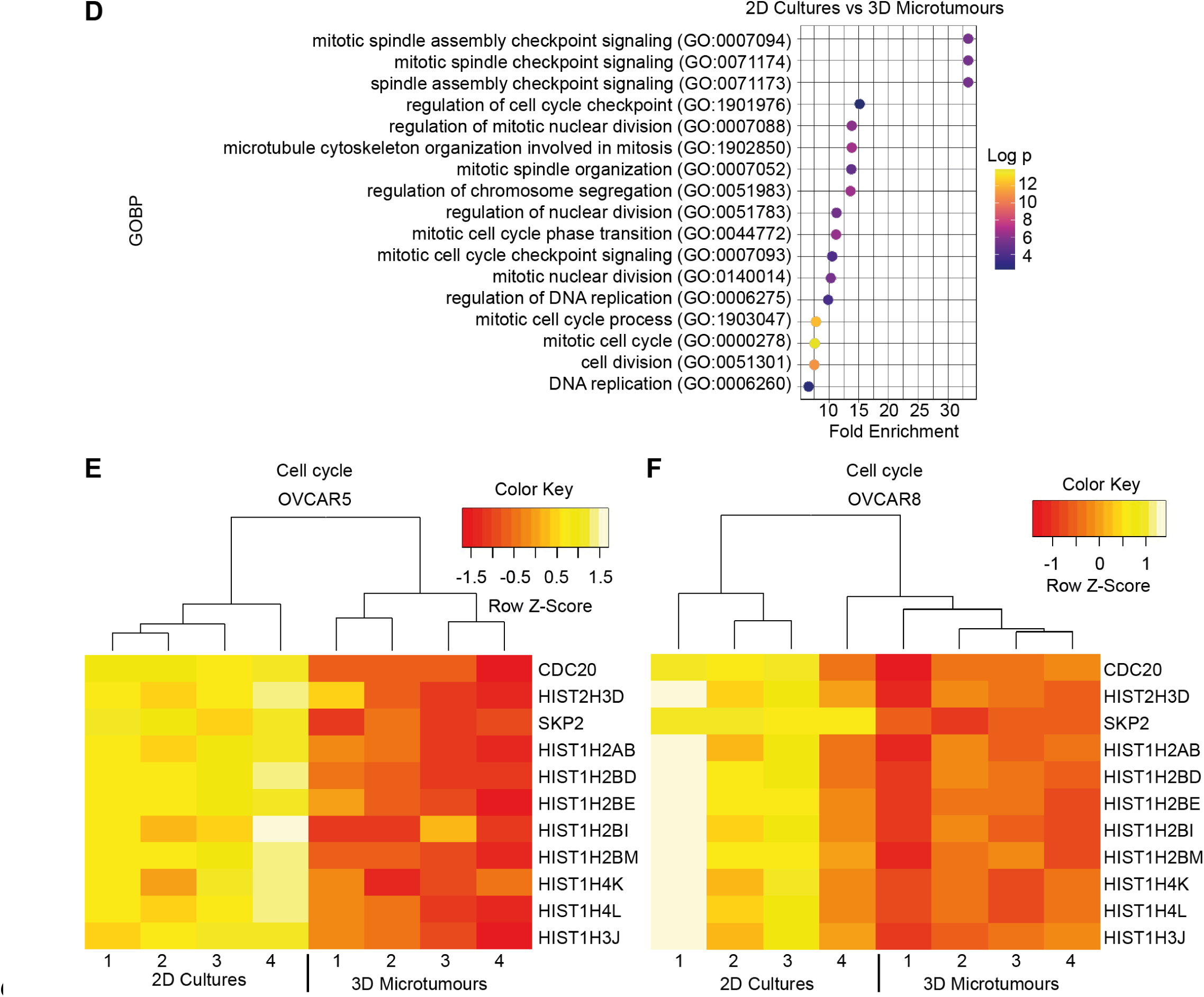
Transcriptomic analysis of 2D cultures and 3D microtumours. Dot plots showing the main biological processes enriched in A) 3D microtumours and D) 2D cultures produced from OVCAR5, OVCAR8 and OVCAR-5/RFP cells. Heatmaps showing the expression of an ovarian cancer specific hypoxic signature in B) OVCAR5 and C) OVCAR-5/RFP. Heatmaps showing the expression of cell cycle related genes in E) OVCAR5 and F) OVCAR8.

On the other hand, 2D cultures were enriched for cell cycle related genes (Figure 3D, E, F), consistent with the faster cell division warranted by the continuous supply of nutrients and oxygen in monolayer systems.

As a whole these results demonstrate that our 3D microtumour system recapitulates key physiological features of tumours that profoundly influence response to therapeutics, and therefore should be a suitable model for drug screening.

### Responses of 3D microtumours to chemotherapy agents

3D microtumours (dimension around 900 µm) composed of ovarian cancer cells were treated with the first-line chemotherapeutics carboplatin and paclitaxel. Freshly prepared 3D microtumours (and 2D cells as for comparison) were distributed into 96-well plates and cultured for 2 days, and then exposed to serial dilutions of carboplatin or paclitaxel for 4 days (Figure S4). Cell viabilities were measured at the end of the treatments and normalized to the corresponding control groups treated with DMSO.

3D microtumours showed higher resistance compared to 2D cells for both anticancer drugs (Figure 4A, B). The ovarian cancer 3D microtumours exhibited IC_50_ values of 100 ± 12.3 µM (carboplatin) and 5.3 ± 2.0 nM (paclitaxel), which were greater than 60 ± 6.1 µM (carboplatin) and 2.7 ± 0.9 nM (paclitaxel) for 2D cells (Figure 4C, D, Table S5).

**Figure 4.**
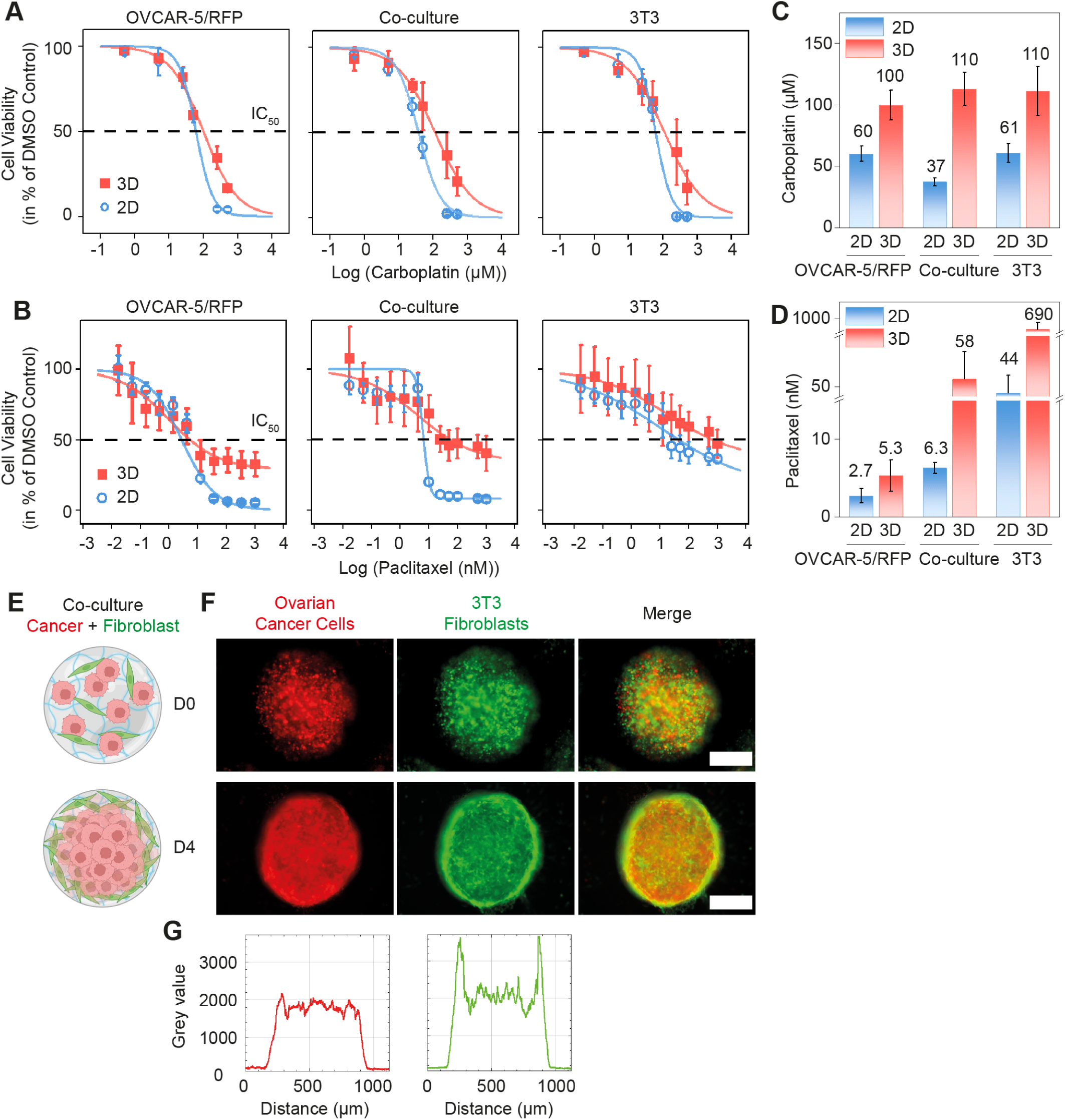
Dose-response of 3D microtumours fabricated by the microfluidic platform. Dose-response curves of 3D microtumours (pink) and 2D cells (blue) treated with serial dilutions of A) carboplatin and B) paclitaxel. Bar graphs of IC_50_ values calculated from the dose-response curves for C) carboplatin and D) paclitaxel. E) Schematic diagram of co-culture 3D microtumours composed of tumour cells and fibroblasts at D0 and D4. The diagrams were created with BioRender.com. F) Epifluorescence images of co-culture 3D microtumours composed of OVCAR-5/RFP and 3T3 fibroblasts on D0 and D4 of fabrication. Scale bar = 300 µm. G) Fluorescence intensity profile of a line across the co-culture 3D microtumour on D4. Left for OVCAR-5/RFP (red) and right for 3T3 fibroblasts (green).

In the clinic, carboplatin and paclitaxel are administered intravenously every three weeks for 8 cycles. Dosage of carboplatin is calculated to deliver an AUC (area under curve) of 5 (mg mL^−1^)⋅min. This would give a theoretical maximum plasma concentration (C_max_) of 280 µM in a typical patient (calculations in Table S6), although the true C_max_ is around 115 µM^34^. *In vivo*, around 50% reduction in the CA125 biomarker is observed with each neoadjuvant chemotherapy cycle^35^. This response aligns to the IC_50_ for carboplatin in 3D microtumours but, by contrast, 2D monolayers are around twice as sensitive. With this said, recapitulating pharmacokinetics is difficult; the *in vitro* carboplatin dose to deliver the same AUC as *in vivo* would be just 6.5 µM. Moreover, *in vivo* the longevity of plasma paclitaxel concentration above 50 nM is what is typically associated with clinical efficacy – 10-fold higher levels than observed *in vitro*^36^.

Stromal cells, such as fibroblasts, are crucial components in the ovarian cancer microenvironment and can regulate tumour progression^37^. Therefore, we incorporated fibroblasts in the ovarian cancer 3D microtumours (co-culture) to provide a more physiological relevant microenvironment. When treated with paclitaxel, a 11-fold higher IC_50_ value was found in co-culture than in OVCAR-5/RFP tumour cells only 3D microtumours. While for 2D cultures, the IC_50_ was only 2.3-fold higher in co-culture than in tumour cells only (Table S7). These IC_50_ values indicate that crosstalk between fibroblasts and tumour cells was different in 3D and 2D cultures, which may contribute to therapeutic failure. Interestingly, we found more cells survived the paclitaxel treatment in co-culture than in tumour cells only. To further investigate the responses to paclitaxel in co-culture, a third collection of 3D microtumours that contained 3T3 fibroblasts only were fabricated. Despite high concentration of up to 1000 nM was used, 47% cells in 3D microtumour and 36% cells in 2D cultures survived the paclitaxel treatment. The result that 3T3 cells were not completely eliminated can be attributed to the differential cytotoxicity of paclitaxel towards tumour cells over normal cells^38^.

Intriguingly, distinct migration patterns were observed for co-culture 3D microtumours. OVCAR-5/RFP tumour cells and 3T3 fibroblast cells were well-mixed upon fabrication and both cell types were homogeneously dispersed throughout the 3D microtumour on D0. The 3T3 fibroblasts started to migrate towards the periphery on D2 and clearly accumulated at the edge of the structure by D4. Conversely, OVCAR-5/RFP tumour cells remained evenly distributed within the 3D microtumour from D0 to D4 (Figure 4E, F, G). This observed core-shell structure may also contribute to the increased IC_50_ values for co-culture 3D microtumours than those composed of tumour cells only.

### 3D microtumours as a superior model of ovarian cancer MRD

After establishing that our 3D microtumour system is physiologically accurate and technically sound, we sought to test whether it could be used to model a specific cancer in a practical application.

One of the biggest challenges in cancer research is finding new therapeutics that can eradicate chemotherapy-resistant cells. This problem is especially relevant in ovarian cancer, which exhibits a high recurrence rate of >80% within 18 months^39^ due to MRD, drug resistant cells that survive first line treatment and initiate relapse (Figure 5A).

**Figure 5.**
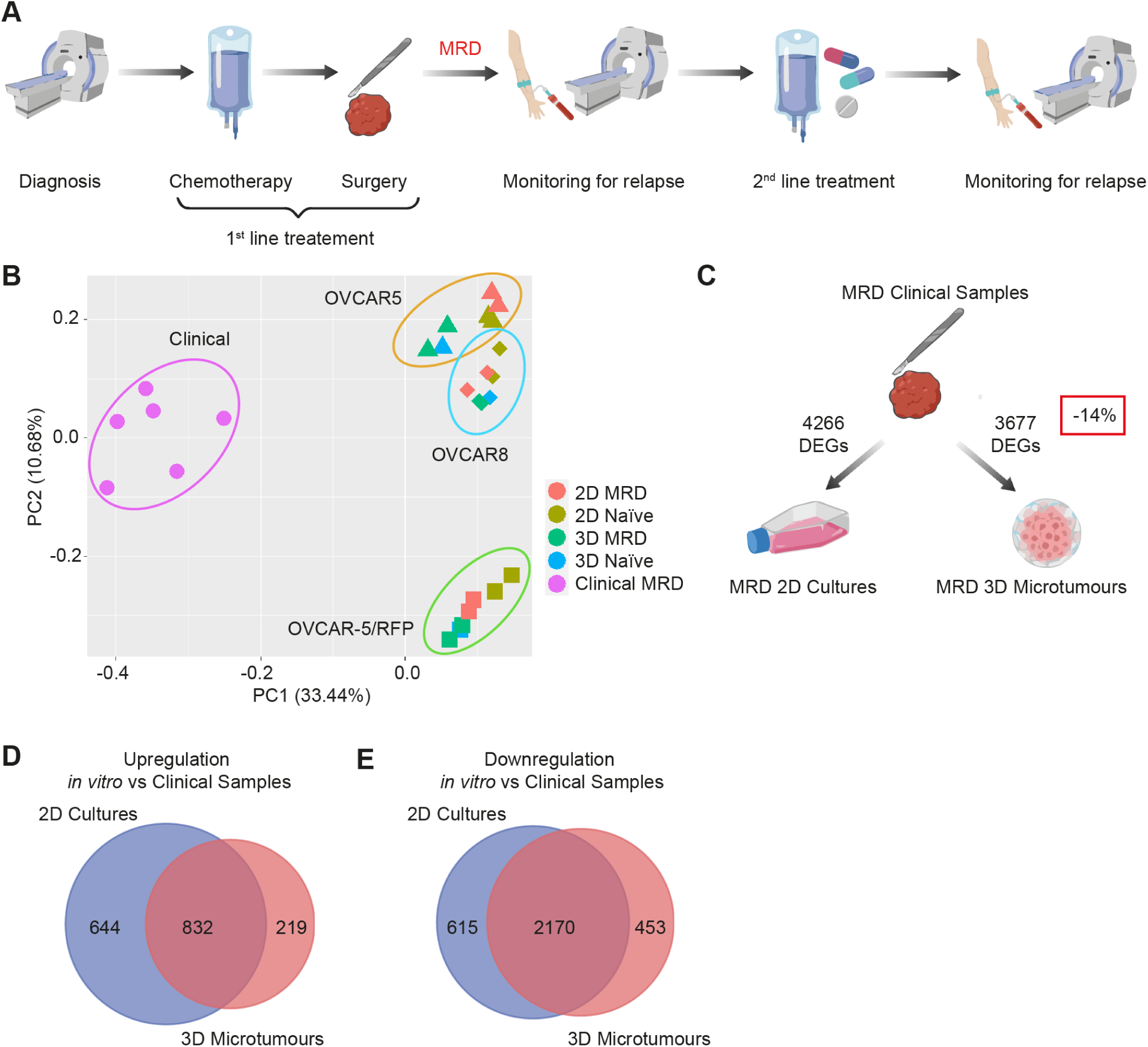
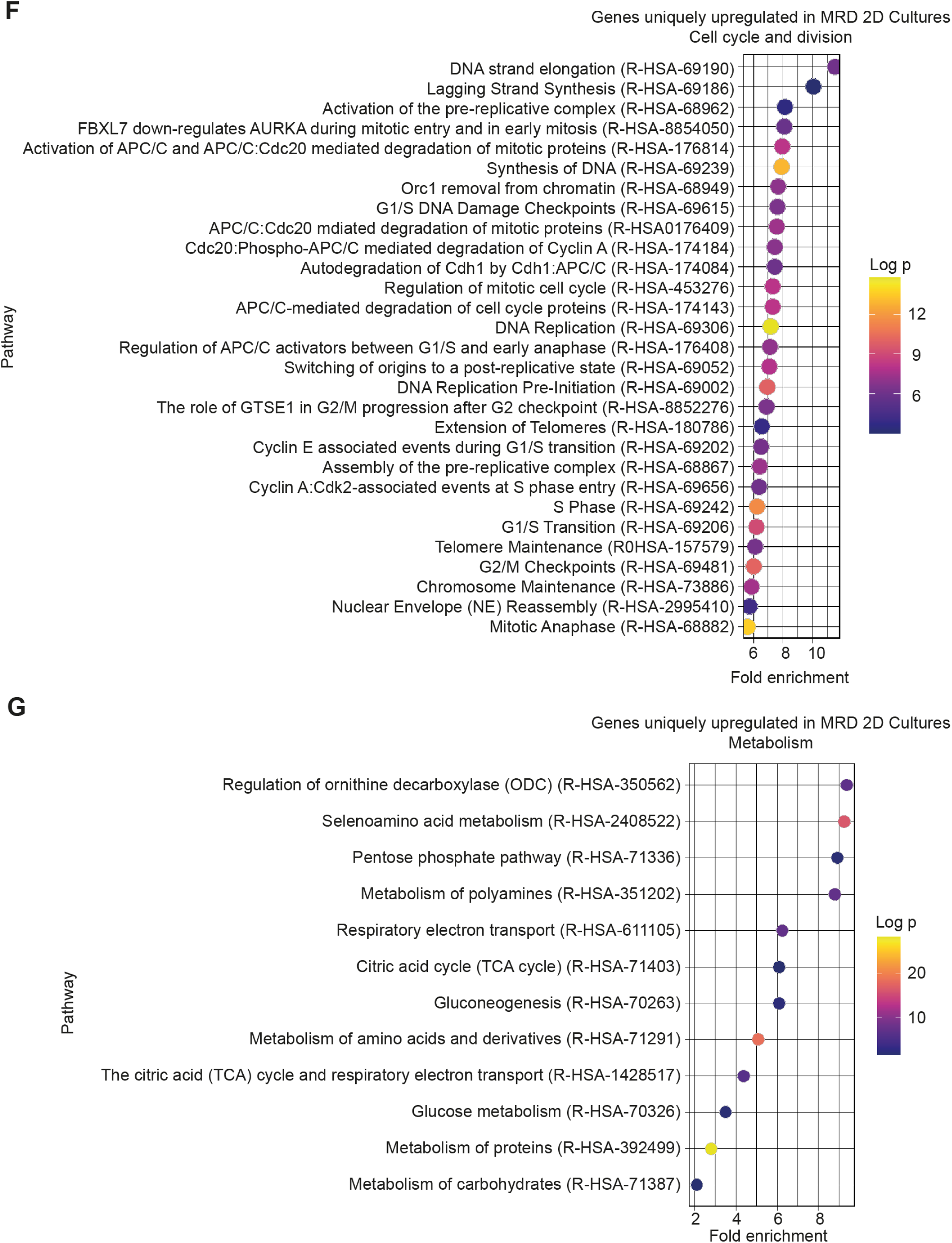
3D microtumours as a superior model of ovarian cancer MRD. A) Schematic diagram of the standard clinical management of patients with ovarian cancer. The diagrams were created with BioRender.com. B) Principal component analysis plots of RNA-Seq data showing 2D cultures, 3D microtumours and the previously published libraries obtained from patients with MRD. C) Number of differentially expressed genes (DEGs) between MRD 2D cultures or MRD 3D microtumours compared to clinical samples. The diagrams were created with BioRender.com. Overlap of DEGs in MRD 2D cultures or MRD 3D microtumours compared to clinical samples for genes that were D) upregulated and E) downregulated *in vitro*. Dot plots showing pathways enriched among genes uniquely upregulated in MRD 2D compared to clinical samples and related to F) cell cycle and division, G) metabolism.

In a previous work we described how MRD cells show distinctive features such as the upregulation of cancer stem cell markers and genes involved in lipid metabolism, and a more pronounced mesenchymal profile^4^. We also developed an MRD 2D *in vitro* model where treatment-naïve cancer cells were exposed to carboplatin concentrations to achieve >90% cell killing; the surviving cells recapitulated some of the features of MRD (such as upregulation of lipid metabolism), but lacked the complexity of multicellularity and three dimensionality. Therefore, we decided to make microtumours from chemotherapy-resistant cells and test their suitability as a model for recapitulating MRD biology.

Firstly, we compared the RNA-Seq data obtained from 2D cultures and 3D microtumours to the previously published libraries obtained from patients with MRD. As shown by Principal Component Analysis (Figure 5B), naïve 2D cells differ most from clinical samples; MRD 2D cells and naïve 3D microtumours represent an intermediate state, while MRD 3D microtumours are the most similar to clinical samples, regardless of the cell line used.

This was confirmed by differential expression analysis, which identified fewer differentially expressed genes (DEGs) between MRD 3D microtumours and clinical samples than between MRD 2D and clinical samples (Figure 5C). The majority of these DEGs are shared between the two comparisons (Figure 5D, E). If we focus only on the genes uniquely upregulated in MRD 2D, we can appreciate a significant enrichment in pathways related to cell cycle and division (Figure 5F), similar to what we observed when we compared the transcriptomes of naïve 2D cultures and naïve microtumours (Figure 3D).

Other genes exclusively enriched in MRD 2D seem to suggest a different metabolic strategy between these cells and the MRD from clinical samples, with the former based on carbohydrates and amino acids (Figure 5G). This was confirmed by the finding that, when we directly compared the transcriptomes of MRD 2D and MRD 3D microtumours, the latter showed significant upregulation of genes involved in lipid transport and metabolism (Figure 6A), similarly to what we originally found in patients with MRD. Moreover, the expression of genes belonging to the original MRD signature correlated significantly with the expression levels observed in the MRD 3D microtumours (Pearson’s correlation coefficient of 0.99, p value <0.05) (Figure S5A, B).

**Figure 6.**
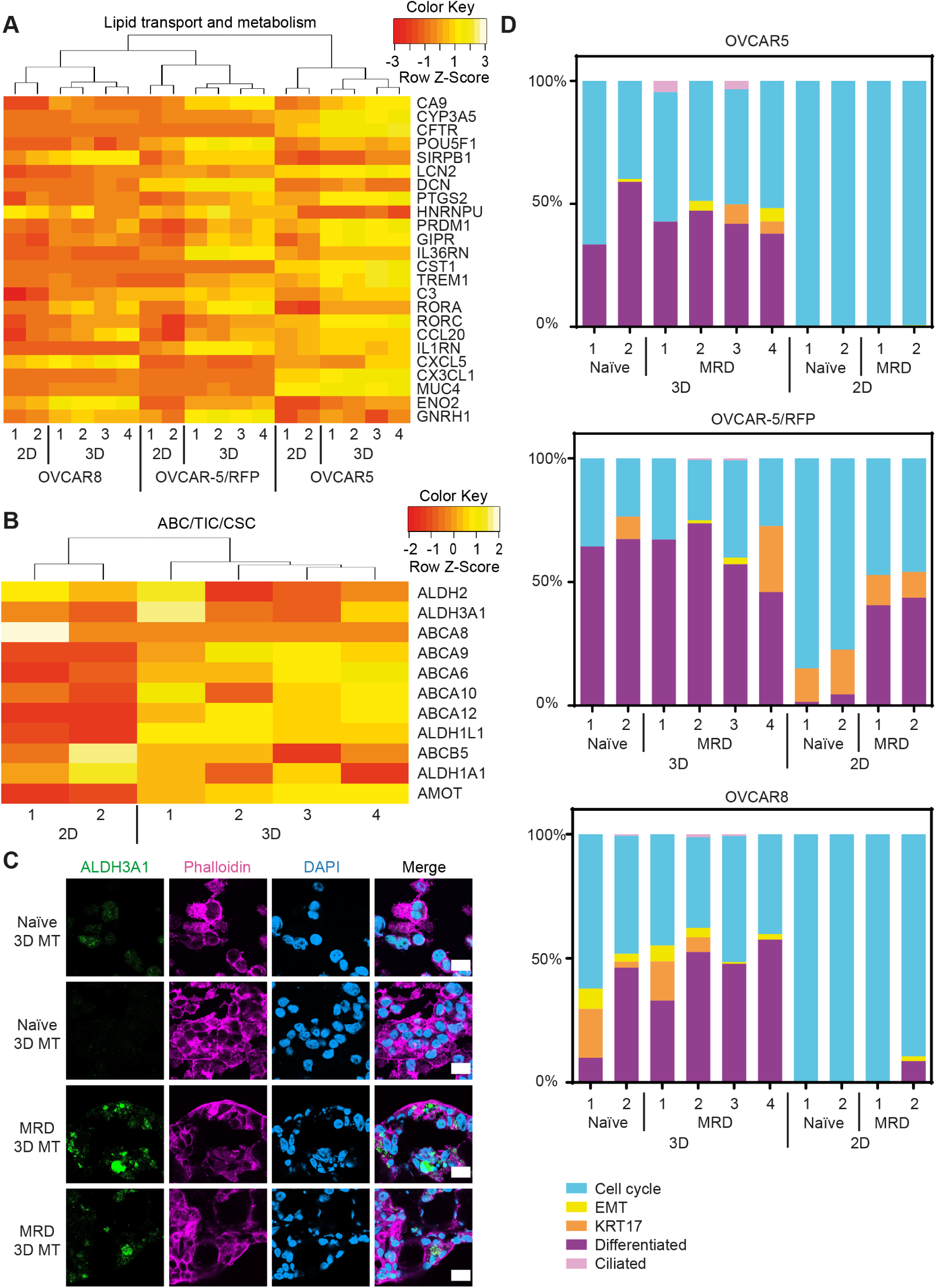
Key features of ovarian cancer MRD recapitulated in 3D microtumours. Heatmap showing the expression of genes in 3D microtumours and 2D cultures A) involved in lipid transport and metabolism, and B) encoding ABC transporters and markers for tumour initiating cells/cancer stem cells (TICs/CSCs). C) Confocal images of 3D microtumours (3D MT) at D10 stained with the TIC/CSC marker: ALDH3A1 (green), phalloidin (pink), DAPI (blue). Scale bar = 20 µm. D) Stacked bar plots visualizing the deconvolution result of 3D microtumours and 2D cultures produced from OVCAR5, OVCAR-5/RFP and OVCAR8. The y axis represents the percentage of each cell state in a given sample. Colours of the bars denote the 5 cell states as shown in the legend.

Another MRD characteristic which is better recapitulated in 3D microtumours is the increased expression of ABC (ATP-binding cassette) transporters and markers for tumour initiating cells/cancer stem cells (TICs/CSCs): consistent with the fact that MRD lesions survive chemotherapy and are the source of ovarian cancer recurrences, we previously identified an ABC/TIC/CSC gene signature which we now find overexpressed in MRD 3D microtumours compared to MRD 2D (Figure 6B). Additionally, we showed that the aldehyde dehydrogenase ALDH3A1, a known TIC/CSC marker which is also important for fatty acid oxidation (FAO)^40^, is specifically expressed in the MRD 3D microtumours but not in the naïve 3D microtumours (Figure 6C).

To further explore the nature of pathways and processes that characterise MRD cells in our different models, we also conducted gene set enrichment analyses (GSEA). This identified several pathways consistently downregulated in both 2D and 3D microtumours relative to clinical samples, such as immunoregulatory interactions (Figure S6A, B) and genes sets associated with ECM (Figure S6C, D). While the downregulation of immunoregulatory interactions is to be expected in our *in vitro* models, both of which lack immune cells, it is important to note the different normalised enrichment score (NES) and adjusted p values for ECM related genes in the two comparisons, pointing to a much more extensive downregulation in the 2D system (Figure S6E). Moreover, the GSEA results confirmed that several cell cycle and non-lipid metabolic pathways are more significantly upregulated in 2D (Figure S6F).

Consistent with this, a recent comparison of the ovarian cancer cell line OVCAR8 in scaffolded spheroids versus monolayers found that the most differentially expressed genes orchestrated immune response, ECM interaction and lipid metabolism^41^.

Finally, we examined whether microtumours can recapitulate non-genetic heterogeneity, a key mechanism for the evolution and survival of cancer cells. Tumour heterogeneity is both genetic and non-genetic, with the latter used to describe cells of the same genetic background but with different phenotypic cell states that can enable invasion, metastasis and chemotherapy resistance.

We previously reported that ovarian cancer non-genetic heterogeneity can be measured with molecular signatures related to five different cell states (cell cycle, EMT (epithelial-mesenchymal transition), KRT17, differentiated and ciliated)^42^. Deconvolution analysis of our RNA-Seq dataset showed that all the five gene signatures originally identified in ovarian cancer clinical samples can be found in the 3D microtumours; however, depending on the cell line, only one to three are present in naïve 2D cultures and, consistent with the results we have shown so far, the most abundant is related to the cell cycle state (Figure 6D). We also analysed a publicly available dataset of 37 additional ovarian cancer cell lines grown in 2D cultures^43^, in all of which the cell cycle signature is dominant if not exclusive (Figure S7A); hence, this is a ubiquitous drawback of monolayer cultures, which all fail to recapitulate the essential features of chemoresistant cells.

On the other hand, 3D microtumours made from cell lines perform at least as well as organoids established from clinical samples^44^, where we observe the occasional sample with only the cell cycle status and very low representation of the ciliated signature (Figure S7B). Furthermore, in our OVCAR5 and OVCAR-5/RFP 3D microtumours, the MRD samples show a higher EMT proportion than the naïve cells (Figure 6D); this is again similar to what we observed in our original characterisation of MRD clinical samples^4^.

Taken all together, these data provide strong evidence in support of using microtumours to model ovarian cancer MRD; the system successfully recapitulates most of its key features, from lipid metabolism to TICs and EMT.

### Using 3D microtumours as a drug screening platform for ovarian cancer MRD

3D microtumours satisfy all the technical criteria to be used as a high-throughput drug screening platform and they can be considered a faithful model for ovarian cancer MRD. The next logical step was to use this system to screen for compounds that can selectively target and eradicate MRD cells.

In our previous work we showed that not only do the MRD cells significantly upregulate their lipid metabolism, but also that this is a vulnerability that can be targeted therapeutically by inhibiting FAO and, in particular, by targeting the carnitine palmitoyl transferase (CPT1) that imports FA into mitochondria for β-oxidation. This was achieved using our 2D model, where MRD cells treated with the CPT1 inhibitors etomoxir and perhexiline underwent 20-30% more cell death than naïve cells^4^.

To compare the previous 2D culture data with our MRD 3D model, 3D microtumours were prepared from naïve and MRD cells and treated with etomoxir and perhexiline for a period of 10 days (Figure S8A). By contrast with the results from 2D cultures, etomoxir failed to induce significant cell death in either naïve or MRD 3D microtumours (Figure 7A), while perhexiline led to a more pronounced reduction in MRD cells in 3D microtumours (Figure 7B) than in 2D cultures. Specifically, in 3D microtumours, perhexiline killed MRD cells 48-82% more effectively than naïve cells, depending on the cell line (Table S8).

**Figure 7.**
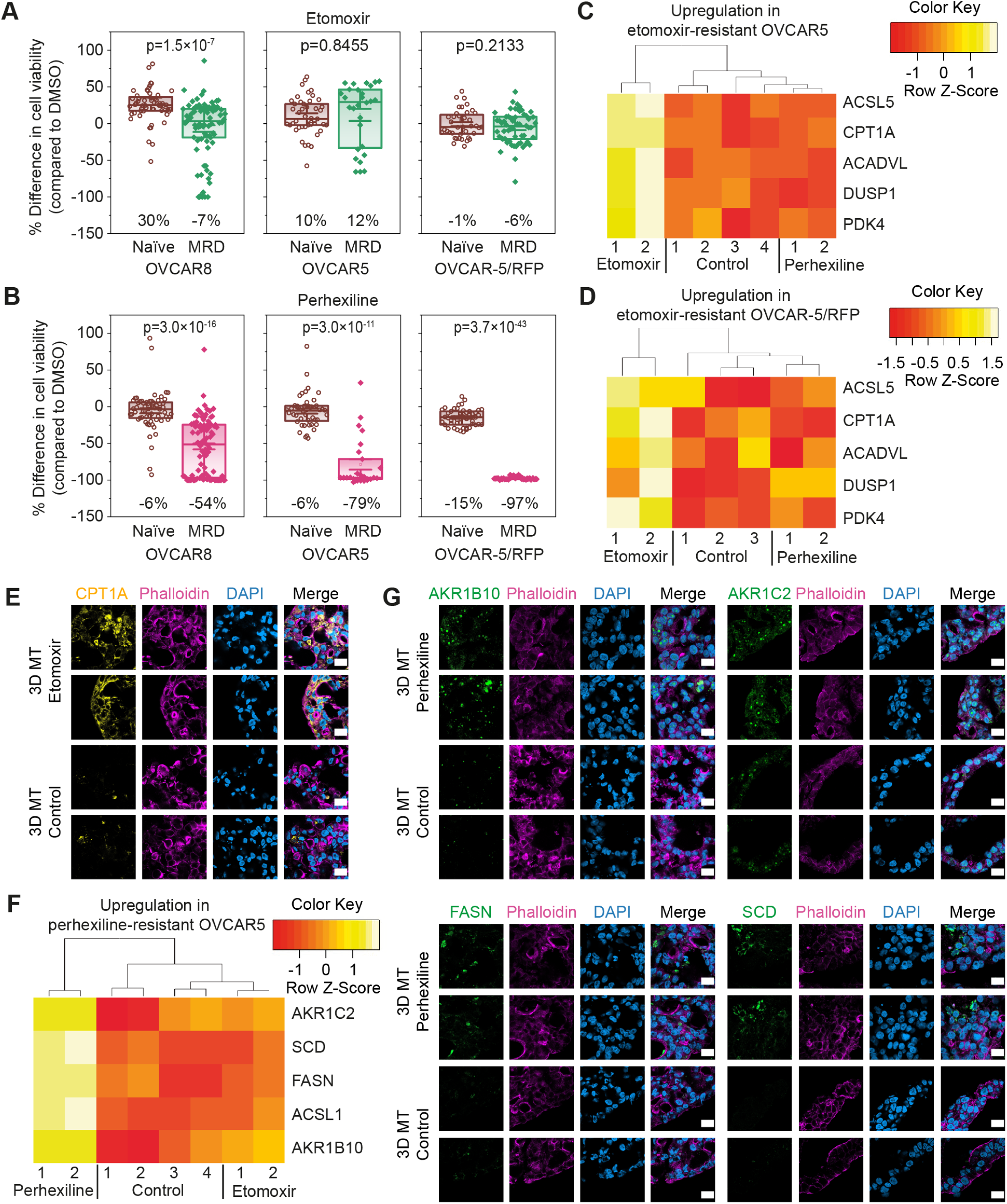
3D microtumours as a drug screening platform for ovarian cancer MRD. Cell viability changes of naïve and MRD 3D microtumours compared to a DMSO control after a 10 d treatment with A) 40 µM etomoxir; B) 4 µM perhexiline (n = 25 to 93). Heatmaps of differentially expressed genes (DEGs) upregulated in etomoxir-resistant MRD 3D microtumours composed of C) OVCAR5 and D) OVCAR-5/RFP. E) Confocal images of 3D microtumours (3D MT) at D10 stained with the FAO marker CPT1A (yellow), phalloidin (pink), DAPI (blue). Scale bar = 20 µm. F) Heatmap of DEGs upregulated in perhexiline-resistant MRD 3D microtumours composed of OVCAR5. G) Confocal images showing the DEGs upregulated in perhexiline-resistant MRD 3D microtumours composed of OVCAR5 at day 10: AKR1B10 (green, top left panel), AKR1C2 (green, top right panel), FASN (green, bottom left panel), SCD (green, bottom right panel), phalloidin (pink), DAPI (blue). Scale bar = 20 µm.

This differential efficacy of FAO inhibitors in 2D cultures versus 3D microtumours could be due to differences in the ability of these compounds to reach the inner cell layers of the 3D microtumours or to their different selectivity (perhexiline inhibits both CPT1 and CPT2 while etomoxir only targets CPT1)^45^.

To gain more insight into this, we analysed the transcriptomes of the cells in the MRD 3D microtumours that survived the treatment with the inhibitors and compared them to the transcriptomes of the cells in the untreated/DMSO control MRD 3D microtumours. Differential analysis showed that, regardless of the cell line, the etomoxir-resistant cells upregulated a set of key genes that can increase FAO at different levels (Figure 7C, D, S8B): beside CPT1 itself, we also found transcripts encoding the long-chain-fatty-acid-CoA ligase ACSL5, the very long-chain specific acyl-CoA dehydrogenase ACADVL, and the pyruvate dehydrogenase kinase PDK4^46^. CPT1 and ACADVL upregulation in the etomoxir-resistant 3D microtumours was also confirmed at protein level through immunofluorescence (Figure 7E, S9).

Extracting enough RNA to make libraries from the perhexiline-resistant 3D microtumours was particularly challenging due to this inhibitor’s strong cytotoxicity and therefore the much lower number of viable cells. However, once we succeeded, the RNA-Seq analysis showed a different gene signature from the etomoxir-resistant 3D microtumours, even though it was still closely linked to lipid metabolism. Depending on the cell line, the cells that survived perhexiline treatment upregulated at least three to five of the following genes at both the RNA and protein level: the long-chain-fatty-acid-CoA ligase ACSL1, the fatty acid synthase FASN, the stearoyl-CoA desaturase SCD and the aldo–keto reductases AKR1B10 and AKR1C2 (Figure 7F, G, S8C, D). While AKR1B10 has been shown to increase FAO in metastatic breast cancer^47^, the inhibition of AKR1C1/2 can sensitise platinum-resistant ovarian cancer towards carboplatin^48^.

Overall, these data confirm once more the potential of the microtumour system, which can be used not only for drug screening purposes but also to investigate resistance mechanisms. More specifically, in this case, our model has enabled us to identify a very promising CPT1/2 inhibitor for targeting ovarian cancer MRD as well as the key genes that we could also target simultaneously to avoid the development of resistance.

## Discussion

While prevention and early detection strategies are essential to improve the survival of cancer patients, there is also a dire need for new therapeutics to successfully eradicate resistant cells. In particular, drugs that could specifically target MRD would be extremely beneficial for women with ovarian cancer, a malignancy with a ten-year survival rate of only 35%^49^, much lower than the 54% for all cancers combined^50^.

In this work we describe how we have achieved the first 3D model of ovarian cancer MRD using microtumours obtained by microfluidics. 3D microtumour models have gained increasing significance over the past decades^51^. Further, the FDA Modernization Act 2.0 of December 2022 has eliminated the requirement for animal tests prior to clinical trials strengthening the position of alternative approaches to drug evaluation^52^. This regulatory change signifies a turning away from animal experiments to more effective models for drug evaluation, as over 90% of drugs reaching the bedside have failed due to efficacy and safety issues^53, 54^.

Here we have showed that our 3D microtumour system represents one such model which satisfies all the technical and biological accuracy requirements to be successfully used in high-throughput drug screening. From a technical point of view, the most important criteria are: 1) uniform size; 2) uniform composition; 3) rapid fabrication; 4) ease of scalability. While uniform size and composition facilitate the comparison of technical repeats, rapid fabrication reduces variations arising from factors such as initial cell population differences. For example, even when cell populations are nearly equalized on D0, the growth rate for each spheroid may differ, leading to variability in screening results. Rapid fabrication and ease of scalability, of course, increase throughput. The 3D microtumours produced by the microfluidic platform meet all the aforementioned requirements, overcoming the weaknesses of conventional spheroids fabrication techniques.

In terms of biological accuracy, our system is able to promptly recapitulate key physiological features observed *in vivo*. Firstly, 3D microtumours exhibiting hypoxia can be generated within 1 day of fabrication (11-21 days for other methods^27, 29^), considerably shortening the time required for high throughput assays. Secondly, the model allows an accurate representation of non-genetic heterogeneity, which is now recognized to play an important role in chemotherapeutic resistance^55^; more specifically, our ovarian cancer 3D microtumours displayed all the molecular signatures related to the five different cell states recently described as the “Oxford Classic”^42, 56^ and which is paving the way to patient stratification for this malignancy.

Lastly, due to their ability to incorporate multiple cell types, the 3D microtumours can recreate the crucial interactions between tumour cells and the microenvironment. This feature cannot be recapitulated in otherwise powerful 3D cultures, like patient-derived organoids, and represents one of their main disadvantages^57^. As an example, we were able to use our model to co-culture ovarian cancer cells and fibroblasts, and we observed the formation of a core-shell structure with fibroblasts migrating and accumulating at the periphery, while the cancer cells did not exhibit a preferred location, a phenomenon that can be attributed to the natural difference in migration ability of the cell types^58^. Ovarian cancer spheroids are known to increase density by reprogramming shell cells to myofibroblasts^59^. This is a form of EMT, and contributes to drug resistance. Our data indicate that the recruitment of local fibroblasts may be an alternative strategy. Moreover, by incorporating different combinations of additional cell species and adjusting their ratios, more complex interactions can be explored with potential therapeutic value. For instance, co-cultures of patient-derived ovarian cancer tumour fragments and autologous immune cells have been shown to enable personalisation of immune checkpoint inhibitor therapy^60^.

Future studies should consider adding other components of the tumour microenvironment, such as adipocytes, to the 3D structure: this is because ovarian cancer is characterised by a very distinctive organ tropism and very rarely spreads outside of the abdominal cavity; one of its preferential metastatic sites is the omentum, where crosstalk between adipocytes and cancer cells has been widely documented^61^. In 2D models of ovarian cancer, adipocyte co-culture confers proliferative and migratory advantage, as well as resistance to cell stress^62^.

In their current composition, our 3D microtumours are already a superior and reliable *in vitro* model for ovarian cancer MRD. Transcriptomics analysis showed that the main aspects of MRD biology related to lipid metabolism and TICs are successfully recapitulated in this system, while the 2D model is limited by the continuous exponential growth of its entire cell population.

Inhibitors of FAO, such as etomoxir and perhexiline, were previously found to selectively kill MRD cells in a 2D format, but to be ineffective towards naïve ovarian cancer cells^4^. When the same treatment scheme was applied to 3D microtumours, surprisingly, etomoxir showed no cell killing while perhexiline had a far greater cytotoxicity towards MRD cells in the 3D than the 2D format. Importantly, the doses of perhexiline at which responses were observed could potentially be delivered safely and locally in ovarian cancer patients using Hyperthermic Intraperitoneal Chemotherapy (HIPEC). Moreover, based on our transcriptomic analysis of the very few cells surviving perhexiline treatment, this population could potentially be eliminated using aldose reductase inhibitors, some of which have successfully been used to reverse drug resistance in prostate and colorectal cancer lines^63^.

## Conclusion

In this study we have developed a 3D model that faithfully recapitulates the characteristics of clinical MRD and can be fabricated *in vitro* in a simple and efficient manner.

As a whole, our findings could lead us a step closer to personalised medicine in the treatment of ovarian cancer. We can imagine a future scenario where biopsies are collected during the diagnostic laparoscopy or following neoadjuvant chemotherapy and then used to create 3D microtumours for the screening of different lipid metabolism inhibitors as well as resistance mechanisms. Through careful regulation of microtumour size and compactness, possible in this model by altering flowrate, tubing size, and cellular co-culture, drug doses could be optimised. Each patient would then receive the drug cocktail that proved to be the most efficient at killing her tumour cells *ex vivo*.

Importantly, this work also represents a crucial proof of concept for the use of 3D microtumours produced by microfluidics as a drug screening platform and, given its versatility, the system could potentially be applied to several different types of tumours.

## Abbreviations

2D: Two-dimensional
3D: Three-dimensional
ABC: ATP-binding cassette
ATCC: American Type Culture Collection
AUC: Area under curve
BSA: Bovine serum albumin
CPT: Carnitine palmitoyl transferase
CSCs: Cancer stem cells
DAPI: 4′,6-diamidino-2-phenylindole
DEGs: Differentially expressed genes
DMEM: Dulbecco’s modified Eagle’s medium
DMSO: Dimethyl sulfoxide
ECM: Extracellular matrix
EMT: Epithelial-mesenchymal transition
FA: Fatty acid
FAO: Fatty acid oxidation
FBS: Fetal bovine serum
FDA: Food and Drug Administration
GSEA: Gene set enrichment analyses
HGSOC: High Grade Serous Ovarian Cancer
HIPEC: Hyperthermic Intraperitoneal Chemotherapy
ID: Inner diameter
JCRB: Japanese Collection of Research Biosources
MRD: Minimal Residual Disease
NEAA: Non-Essential Amino Acids
NES: Normalised enrichment score
OCT: Optimal cutting temperature
PBS: Phosphate-buffered saline
PDMS: Polydimethylsiloxane
Pen-Strep: Penicillin-Streptomycin
PFA: Paraformaldehyde
PTFE: Polytetrafluoroethylene
RT: Room temperature
TICs: Tumour initiating cells

## Acknowledgements

Work in the HB laboratory was supported by a European Research Council Advanced Grant (SYNTISU) and a Proof of Concept Grant (BIOELECTRIC), and the Oxford Martin School Programme on 3D Printing for Brain Repair, and a Cancer Research UK’s Pioneer Award Grant (CRUK). Work in the AAA laboratory was funded by Ovarian Cancer Action, the Cancer Research UK Oxford Centre and the Medical and Life Sciences Translational Fund of the University of Oxford. R.K.K. was funded by the Health Research Bridging Salary Scheme (0011044) at the University of Oxford. The authors extend their appreciation to the Deputyship for Research & Innovation, Ministry of Education in Saudi Arabia for funding this research work through the project number 852.

## Methods

### Cell Culture

OVCAR5/RFP, NIH3T3/GFP, MDA-MB-231/RFP and HeLa/GFP cell lines were purchased from Cell Biolabs Inc., USA. HEK293T and 3T3-L1 cell lines were purchased from ATCC. Kuramochi cells were obtained from the JCRB Cell Bank. All cells except 3T3-L1 were cultured in DMEM (Sigma-Aldrich, #D5796), supplemented with 10% (v/v) FBS (Sigma-Aldrich, #F7524), 2 mM GlutaMAX™ Supplement (Gibco, #35050038), 0.1 mM MEM NEAA (Sigma-Aldrich, #M7145), and 1% (v/v) Penicillin-Streptomycin (Pen-Strep, 100 U mL^−1^ and 100 μg mL^−1^ respectively, Sigma-Aldrich, #P4333). 3T3-L1 cells were cultured routinely in DMEM supplemented with 10% (v/v) bovine calf serum (ATCC, #30-2020) and 1% (v/v) Pen-Strep. To induce differentiation of 3T3-L1 cells into adipocytes, the cells were cultured in differentiation medium following an established protocol (Table S9)^64^.

### Bioink Preparation

Matrigel® Matrix (#354234) and Collagen I (#354236) were purchased from Corning Life Sciences, UK. The gels were thawed completely on ice before use. Collagen I solution (2 mg mL^−1^) was prepared by diluting Collagen I (3.78 mg mL^−1^, 52.9 μL) with ice-cold DI-water (39.5 μL), 10X DPBS (6.15 μL) and 1 N NaOH (1.2 μL). Agarose solution (2% w/v) was prepared by dissolving agarose powder (Thermo Fisher, #16520050) in sterile water at 100°C, then cooled to 37°C. Silk fibroin solution (50 mg mL^−1^, Sigma-Aldrich, #5154) was thawed at 4°C and supplemented with 10 U mL^−1^ horseradish peroxidase (type VI lyophilized powder, Sigma-Aldrich) and 0.4 μL mL^−1^ hydrogen peroxide solution (30% w/w, Sigma Aldrich). The bioink, with cell density = 3 to 4 x 10^7^ cells mL^−1^, was prepared by resuspending cell pellets in the desired pre-gel solution (Table S1).

### Microfluidics Platform and 3D Microtumour Fabrication

The microfluidics platform was improved over work previously reported by our group^65^. The PDMS microfluidic chips (Figure S1A) were prepared by casting on a custom-made reverse mould, which was produced by a three-dimensional printer (Solid Print3D, Formlabs) using clear resin (Formlabs), and provided more choices of channel size compared to the previous method of drilling holes in PDMS blocks to make the T-junction. The cell-laden bioink and the oil, tetradecane (Sigma-Aldrich, #172456), were loaded into separate syringes (Figure 1), and pumped into the 3-channel microfluidic chip with neMESYS syringe pumps (Cetoni, Korbussen, Germany). Droplets containing cells in Matrigel, separated by the oil, were formed in a PTFE exit tube (Cole-Parmer, UK). Upon complete gelation (Table S1), the 3D microtumours were ejected from the exit tube, transferred to cell culture medium and maintained at 37°C, 5% CO_2_. Co-culture 3D microtumours in this work composed of a 50: 50 mixture of OVCAR-5/RFP tumour cells and 3T3 fibroblasts.

### Characterizations of 3D Microtumours

#### Size Distribution

The 3D microtumours were imaged by using a Leica DMi8 inverted epifluorescence microscope platform equipped with a Leica DFC7000 CCD camera (Leica Microsystems Ltd, UK). Images were processed with Fiji ImageJ software to obtain the diameter of each 3D microtumour. For 3D microtumours with the cross-section of an ellipse, the dimensions were defined as 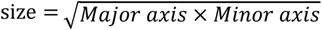.

#### Cell Viability

The viabilities of cells in 2D culture and bioinks were determined with a Countess™ Automated Cell Counter (Invitrogen) by using 0.4% trypan blue solution (Invitrogen, #C10314).

The viabilities of cells in 3D microtumours were evaluated with PrestoBlue™ Cell Viability Reagent (Thermo Fisher, #A13261) according to the manufacturer’s instructions and fluorescence intensity was measured with a microplate reader (CLARIOstar Plus, BMG LABTECH) (Table S10).

#### Hypoxic Core Staining

Image-iT™ Green Hypoxia Reagent (Thermo Fisher, #I14834) was dissolved in DMSO (Sigma-Aldrich, #D8418) to prepare a 5 mM stock solution, which was added to culture medium at a final concentration of 5 µM. After incubation at 37°C for 3 h, 3D microtumours were imaged with a confocal laser scanning microscope (Leica TCS SP5, Leica Microsystems). A standard FITC/GFP excitation/emission filter set was applied. Z-stack images were taken with an optical section thickness of 10 µm. The optical z slices were projected to form 2D images using z-project in Fiji ImageJ software.

### Anticancer Drug Responses

Carboplatin powder (Cayman Chemical, #13112) was dissolved and then serially diluted with sterile water to yield a range of working solutions, which were added to cell medium at a 1:20 volume ratio to a maximum final concentration of 500 µM.

Paclitaxel powder (Invitrogen, #P3456) was dissolved and then serially diluted with DMSO to yield a range of working solutions, which were added to cell medium at a 1:100 volume ratio to a maximum final concentration of 1 µM.

For drug treatment, one microtumour or 5000 cells (2D) were seeded into each well of a 96-well plate (Corning #3595). After 2 d, the medium was replaced with drug-containing medium, and the treatment was continued for 4 d. PrestoBlue was used to evaluate cell viability at the end of the drug treatment. For each condition, 2D cells n = 11 to 21, 3D microtumours n = 20 to 32.

OriginPro 2023 (OriginLab Corporation) was used to plot cell viability data and generate fitted dose-response curves. The IC_50_ values were derived from the dose-response curves at 50% cell viability.

### Minimal Residual Disease Modelling

#### Preparation of MRD-like cells

OVCAR5 and OVCAR8 cell lines were obtained from ATCC and cultured in RPMI 1640 (Gibco, Thermo Fisher, #21875034), supplemented with 10% (v/v) FBS and 1% (v/v) Pen-Strep.

To prepare MRD-like cells, 2D naïve cancer cells were treated with carboplatin for 2 weeks at specific optimised conditions to achieve more than 90% cell killing as previously described^4^. All the cells collected on D14 were expanded for 2 to 14 days (depending on cell growth) to produce the MRD cells for later use.

#### Fatty Acid Oxidation Inhibitor Responses

Etomoxir sodium salt (Stratech, #S8244-SEL) was dissolved in DMSO to prepare a 40 mM stock solution, which was diluted in cell medium to a final concentration of 40 µM. Perhexiline (Cambridge Bioscience, #CAY16982) was dissolved in DMSO to prepare a 4 mM perhexiline stock solution, which was diluted in cell medium to a final concentration of 4 µM. DMSO was added to cell medium at a 1:1000 volume ratio for the control group.

3D microtumours were prepared from both naïve and MRD cells. Five 3D microtumours were seeded into each well of a 12-well plate on the day of fabrication and cultured with 2 mL of drug-containing or DMSO-containing medium for 10 d. PrestoBlue was used to evaluate cell viability at the end of the drug treatment.

#### RNA Extraction and Library Preparation

RNA was extracted with the RNAqueous-Micro Total RNA Isolation Kit (Thermo Fisher, #AM1931). RNA integrity was evaluated by RIN value with the 2200 TapeStation System (Agilent Technologies, Inc.) and only samples with RIN values above 7 were taken forward for library preparation, which was performed using a KAPA HyperPrep Kit (Kapa Biosystems, #KR1351) following the manufacturer’s instructions. The libraries were evaluated by using the 2200 TapeStation System and then quantified with a Qubit 2.0 Fluorometer (Thermo Fisher, Invitrogen). Multiplexed library pools of different samples were quantified with the KAPA Library Quantification Kit (Roche) and sequenced by using 75 bp paired-end reads on the NextSeq500 platform (Illumina).

#### Processing of RNA-Seq data

Sequencing reads from FASTQ files were trimmed for adapter sequences and quality with Trim Galore!, and mapped to the UCSC hg19 human genome assembly using STAR (v2.7.3a). Read counts were obtained by using subread FeatureCounts (v2.0.0).

Differential expression analysis was carried out by using edgeR (v3.36.0). Statistical overrepresentation analysis was performed with PANTHER (v17), and the threshold for significance was set at FDR < 0.05.

Deconvolution analysis was performed as previously described ^42^ in the relative mode, and thus, for each tumour the scores of the 5 molecular signatures added up to 1.

#### Study approval

The HGSOC clinical samples used in this study were recruited under the Gynaecological Oncology Targeted Therapy Study 01 (GO-Target-01, NHS Health Research Authority South Central – Berkshire Research Ethics Committee research ethics approval 11-SC-0014) and the Oxford Ovarian Cancer Predict Chemotherapy Response Trial (OXO-PCR-01, NHS Health Research Authority South Central – Berkshire Research Ethics Committee research ethics approval 12-SC-0404). All participants involved in this study were appropriately informed and consented.

#### Immunofluorescence staining

The 3D microtumours and the clinical samples were embedded in OCT (NEG-50, Richard-Allan Scientific), frozen and kept at −80°C until sectioning. 10 μm sections were taken in a CryoStar cryostat microtome (Thermo Fisher) and stained for immunofluorescence imaging. The slides were washed with ice-cold PBS twice to remove the OCT, fixed in 4% PFA for 10 min and permeabilized with 0.1%TritonX-100 in PBS for 10 min at RT. The samples were then incubated in Blocking Buffer (2% BSA + 0.1% TritonX-100 in PBS) for 1 hour followed by an overnight incubation with the diluted primary antibodies (Table S11) in a humidified chamber at 4°C. The following day the slides were washed in PBS and incubated with the secondary antibodies and phalloidin for 1 hour at RT. After extensive washes in PBS, the slides were mounted with Vectashield + DAPI (VectorLaboratories) and dried in the dark before being imaged using a confocal microscope (Zeiss900).

## Author contributions

X.Y., M.A., H.B. and A.A.A. conceived and designed the work, which was supervised by H.B. and A.A.A. X.Y. improved the microfluidic platform, and performed 3D microtumour fabrication and chemotherapy assays. X.Y. and M.A. contributed to cell culture and manipulation, microscope imaging, cryo-sectioning, RNA-Seq library preparation and data and image analysis. M.A. performed the bioinformatics analysis and immunofluorescence. L.R. performed the deconvolution analysis. Y.J. and L.Z. contributed to cryo-sectioning and cell staining. Y.Z. contributed to the preparation of silk fibroin and reverse moulds of microfluidic chips. E.M. contributed to cell monolayer culture. N.M. assisted with preliminary experiments on chemotherapeutic screening. R.K.K., S.M., A.Aggarwal and L.Z. contributed discussions and helped with image analysis. A.Albhukari contributed discussions. X.Y., M.A., A.Aggarwal, H.B. and A.A.A. wrote the manuscript. H.B., A.A.A., L.Z. and M.A. acquired funding. All authors read and revised the manuscript.

