## Supplementary for "A 3D microtumour system that faithfully represents ovarian cancer minimal residual disease"

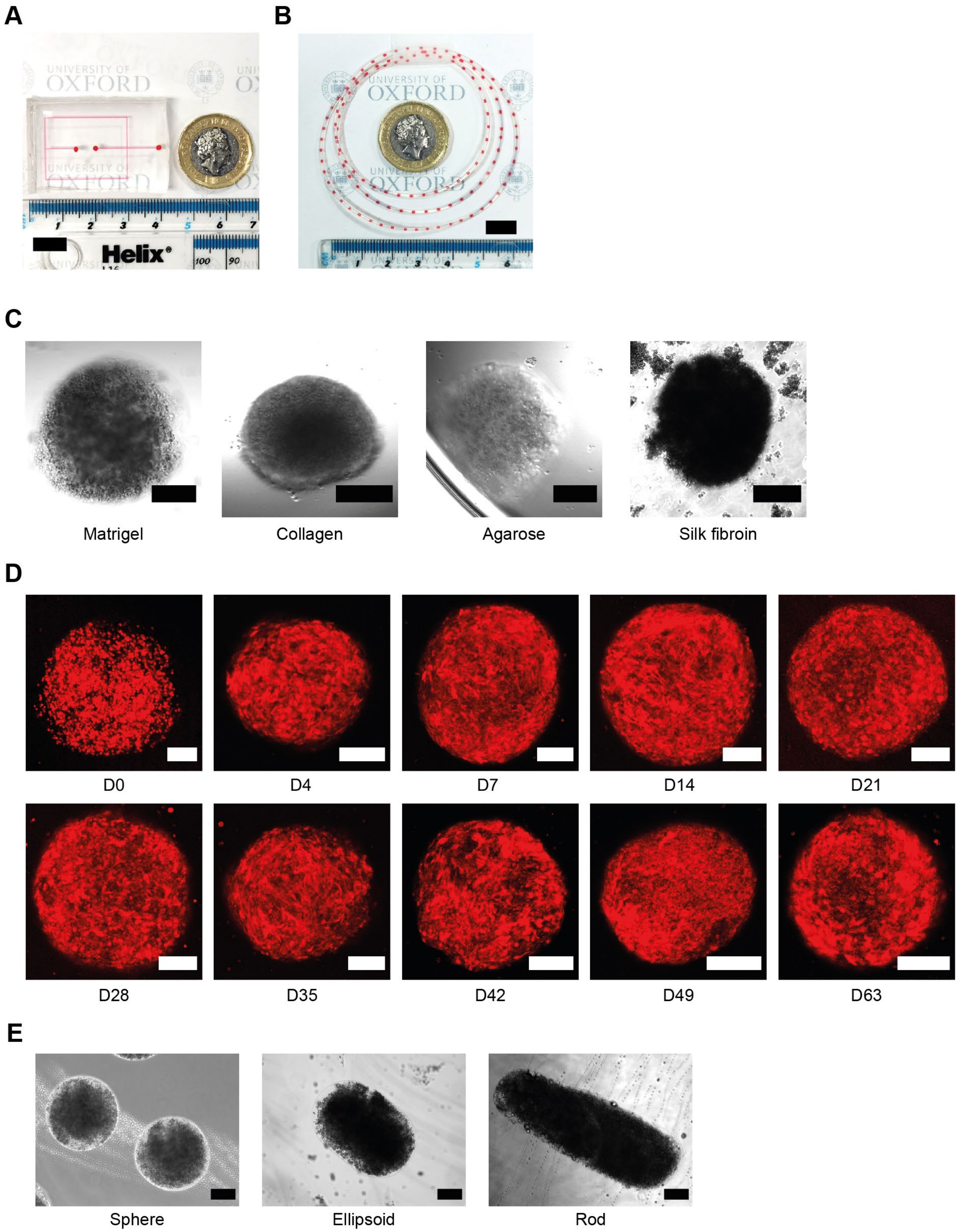


**Figure S1.** A) Image of a 3-channel PDMS microfluidic chip. Scale bar = 1 cm. B) Image of 3D microtumours (stained with red dye) in a PTFE exit tube. Scale bar = 1 cm. C) 3D Microtumours composed of OVCAR8 and different hydrogels. Bright field microscope images for Matrigel, collagen, agarose and silk fibroin microtumours were taken on D0 of fabrication. Scale bar = 300 μm. D) Long term culture of 3D microtumours. The 3D microtumours were composed of OVCAR-5/RFP (red) and Matrigel. Fluorescence microscope images were taken from D0 to D63 after fabrication. Scale bar = 300 μm. E) Different shapes of 3D microtumours. 3D microtumours were composed of OVCAR8 and Matrigel. Bright field microscope images were taken on D0 of fabrication. The shapes of sphere, ellipsoid and rod were achieved by adjusting the bioink:oil flowrate ratio to 1000:3000 μL $h^{-1}$, 1000:1000 μL $h^{-1}$, and 1000:300 μL $h^{-1}$, respectively. Scale bar = 300 μm.

**Table S1.** Summary of biocompatible hydrogels and their preparation conditions used in 3D microtumour fabrication.

| Hydrogels | Preparation for bioink | Microfluidic fabrication temperature | Post-fabrication process temperature | Gelation time |
|---|---|---|---|---|
| Matrigel | Thaw completely on ice, then mix with cells on ice. | 8°C | 37°C | 2 h |
| Collagen | Thaw completely on ice, then mix with cells on ice. | 8°C | 37°C | 1.5 h |
| Agarose | Melt at 70°C and cool to 37°C, then mix with cells at 37°C. | 37°C | 4°C | 10 min |
| Silk fibroin | Thaw completely at 4°C, then mix with cells on ice. | 8°C | 37°C | 30 min |

**Table S2.** Summary of cells used in 3D microtumour fabrication

| Cell line | Classification | Source | Catalogue No. |
|---|---|---|---|
| OVCAR-5/RFP | Ovarian cancer | Human | Cell Biolabs Inc. AKR-254 |
| OVCAR5 | Ovarian cancer | Human | ATCC |
| OVCAR8 | Ovarian cancer | Human | ATCC |
| Kuramochi | Ovarian cancer | Human | JCRB CVCL_1345 |
| MDA-MB-231 | Breast cancer | Human | Cell Biolabs Inc. AKR-251 |
| HeLa/GFP | Cervical carcinoma | Human | Cell Biolabs Inc. AKR-213 |
| NIH3T3/GFP | Fibroblast | Mouse | Cell Biolabs Inc. AKR-214 |
| 3T3-L1 derived adipocytes | Fat cell | Mouse | ATCC CL-173 |
| HEK293T | Embryonic kidney | Human | ATCC CRL-3216 |

**Table S3.** Cell viability data at three stages during the microfluidic fabrication of 3D microtumours from various cells and hydrogels (n = 3 to 6).

| Cell | Hydrogel | Cell viability | | |
|---|---|---|---|---|
| | | 2D cells* | Bioink | 3D microtumours |
| OVCAR8 | Matrigel | 93% ± 2.0% | 95% ± 0.58% | 96% ± 0.74% |
| OVCAR-5/RFP | Matrigel | 93% ± 2.7% | 95% ± 0.85% | 93% ± 2.1% |
| 3T3 | Matrigel | 94% ± 1.9% | 97% ± 0.15% | 93% ± 0.45% |
| OVCAR8 | Agarose | 95% ± 2.0% | 95% ± 0.59% | 93% ± 0.68% |
| OVCAR8 | Collagen | 95% ± 2.0% | 96% ± 1.1% | 97% ± 0.34% |

* 2D cells were harvested from culture flasks.

**Table S4.** Dimensions of 3D microtumours formed from Matrigel and OVCAR-5/RFP tumour cells, 3T3 fibroblast cells and a 50: 50 mixture of OVCAR-5/RFP cells and 3T3 cells (co-culture). Size was measured on D0 of fabrication.

| Cell | Size Group | 3D Microtumour Size (μm) | | | |
|---|---|---|---|---|---|
| | | N total | Median | Mean | Standard Deviation |
| OVCAR-5/RFP | Small | 84 | 321 | 321 ± 7.17 | 2.2% |
| OVCAR-5/RFP | Medium | 77 | 672 | 669 ± 19.8 | 3.0% |
| OVCAR-5/RFP | Large | 90 | 924 | 927 ± 24.7 | 2.7% |
| Co-culture | Small | 88 | 302 | 302 ± 7.49 | 2.5% |
| Co-culture | Medium | 76 | 675 | 683 ± 39.4 | 5.8% |
| Co-culture | Large | 86 | 915 | 920 ± 26.4 | 2.9% |
| 3T3 | Small | 62 | 314 | 317 ± 11.7 | 3.7% |
| 3T3 | Medium | 74 | 699 | 696 ± 22.8 | 3.3% |
| 3T3 | Large | 84 | 994 | 995 ± 28.5 | 2.9% |

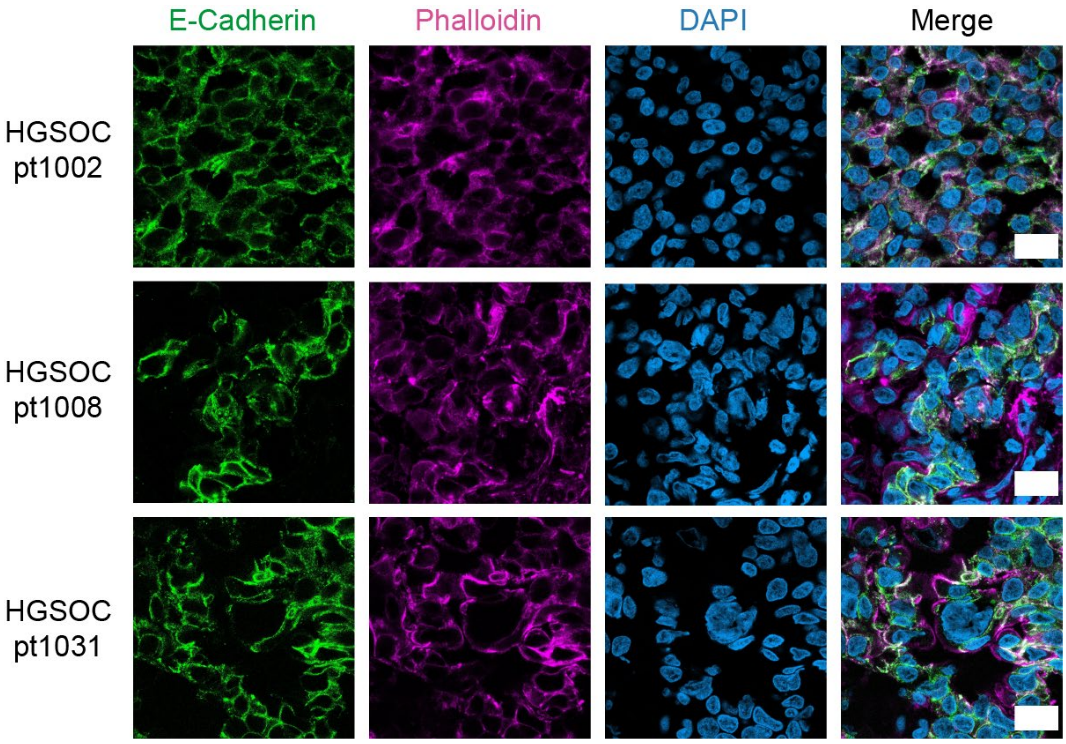


**Figure S2.** Confocal images of clinical samples stained with E-Cadherin (green), phalloidin (pink), DAPI (blue). Scale bar = 20 μm.

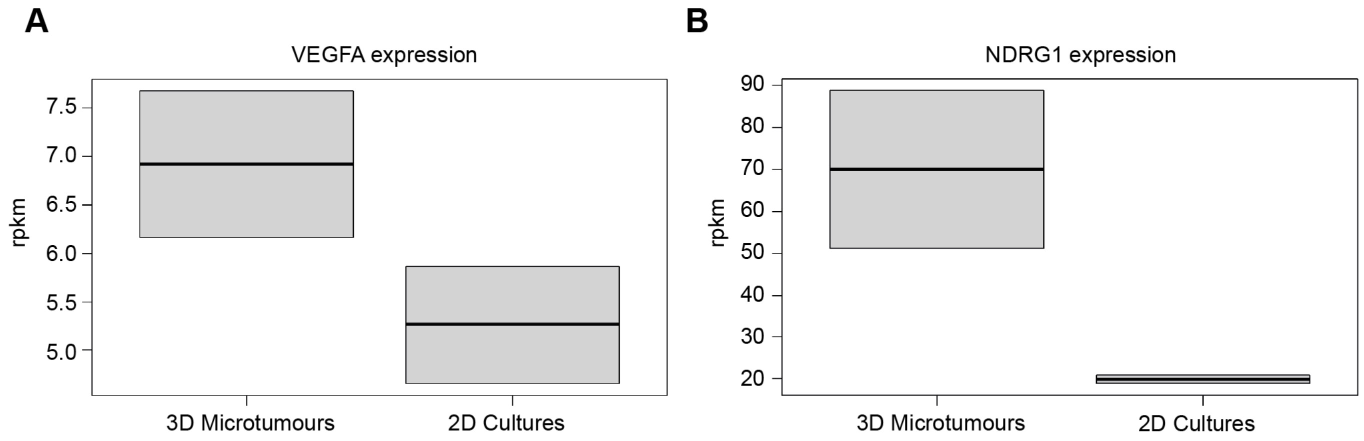


**Figure S3.** Box plots showing the expression of the hypoxia related genes A) VEGFA and B) NDRG1 in OVCAR8 3D microtumours and 2D cultures (rpkm = reads per kilo base per million mapped reads).

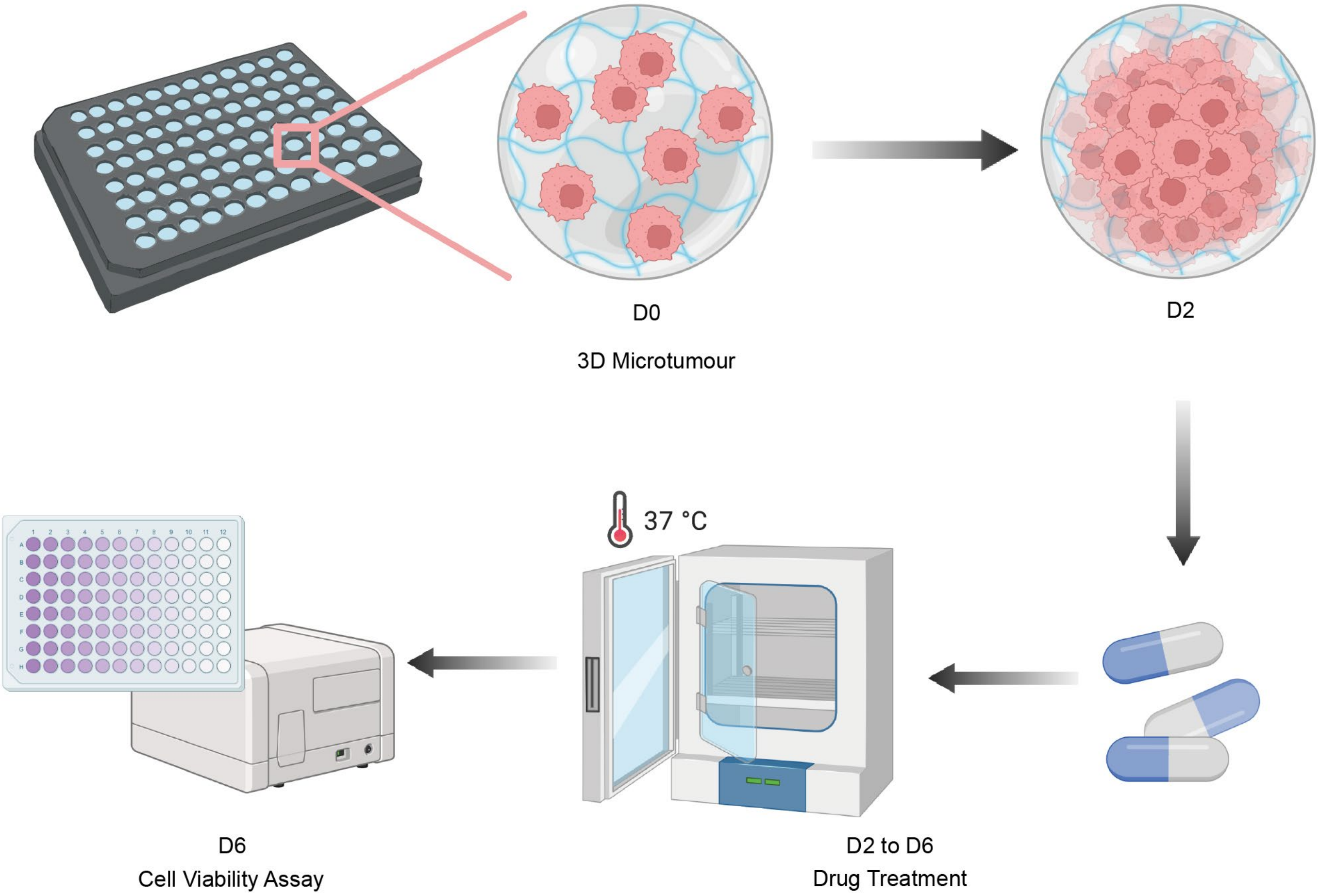


**Figure S4.** Schematic showing the evaluation of chemotherapeutic responses with 3D microtumours. Each 3D microtumour was seeded to one well of a 96-well plate on D0 of fabrication and kept at 37°C for 2 days. On D2, serial dilutions of chemotherapeutics were applied and the microtumours were cultured at 37°C for 4 days. On D6, a cell viability assay was performed for each 3D microtumour. The diagrams were created with BioRender.com.

**Table S5.** $IC_{50}$ values calculated from carboplatin and paclitaxel dose-response curves.

| | $IC_{50}$ | 2D | 3D |
|---|---|---|---|
| OVCAR-5/RFP | Carboplatin (μM) | 60 ± 6.1 | 100 ± 12 |
| | Paclitaxel (nM) | 2.7 ± 0.9 | 5.3 ± 2.0 |
| Co-culture | Carboplatin (μM) | 37 ± 3.1 | 110 ± 14 |
| | Paclitaxel (nM) | 6.3 ± 0.7 | 58 ± 27 |
| 3T3 | Carboplatin (μM) | 61 ± 7.6 | 110 ± 20 |
| | Paclitaxel (nM) | 44 ± 18 | 690 ± 210 |

**Table S6.** Calculations for theoretical carboplatin $C_{max}$ and *in vitro* equivalent dose

| Theoretical carboplatin plasma $C_{max}$: | Equivalent *in vitro* dose for carboplatin AUC: |
|---|---|
| First, consider the average epithelial ovarian cancer patient parameters of age 63, BMI 26, creatinine 0.85 mg $dL^{-1}$ and female gender, and use the Cockcroft-Gault formula to estimate GFR.<br>Next, use the Calvert formula to calculate carboplatin dose: Dose (mg) = AUC (mg $mL^{-1}$)·min x [GFR (mL $min^{-1}$) + 25 (mL $min^{-1}$)]<br>Assume no excretion and serum-only distribution during the 30-minute infusion window, with a typical blood volume of 4.7 L, to yield the theoretical $C_{max}$ of 280 µM. | *In vitro*, we assume no distribution phase, and a prolonged half-life of 24 h.<br>Using $A = A_0e^{-kt}$ with $A/A_0 = ½$ and t = 1440, we obtain k = ln(2)/1440.<br>Integrating $A = A_0e^{-kt}$ between ∞ and 0, and equating the result to an AUC of 5 (mg $mL^{-1}$)·min, we find $A_0$ = 5k= 0.0024 mg $mL^{-1}$ = 6.5 µM. |

**Table S7.** Comparison of paclitaxel $IC_{50}$ values of co-culture over OVCAR-5/RFP.

| Paclitaxel $IC_{50}$ (nM) | Co-culture | OVCAR-5/RFP | Fold change for $IC_{50}$ value of Co-culture over OVCAR-5/RFP |
|---|---|---|---|
| 3D | 58 ± 27 | 5.3 ± 2.0 | 11 |
| 2D | 6.3 ± 0.7 | 2.7 ± 0.9 | 2.3 |

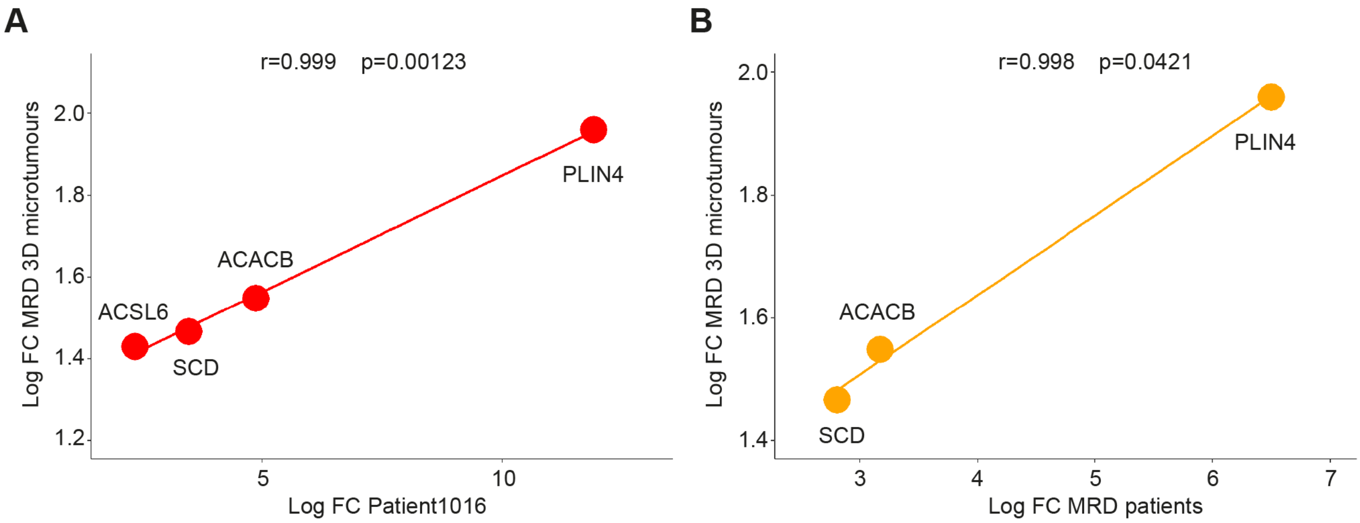


**Figure S5.** Scatterplots showing the $\log_2$ fold change (logFC) of genes involved in lipid metabolism. A positive correlation is observed between their upregulation in the MRD 3D microtumours (3D/2D expression ratios) compared to the MRD patient 1016 (A) or the pooled MRD exceptional responders (B) (post/pre chemo expression ratios).

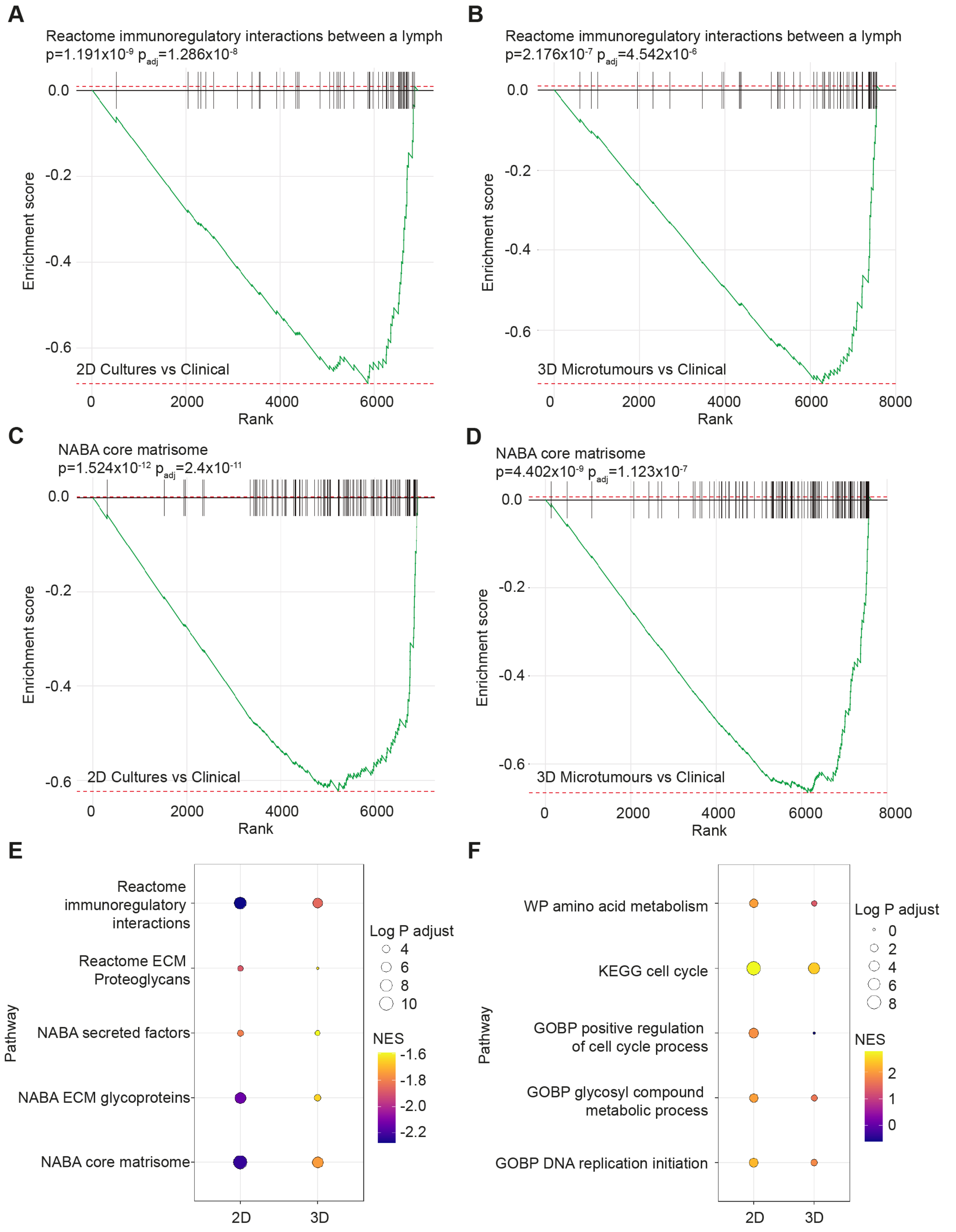


**Figure S6.** Results of gene set enrichment analyses (GSEA) of pathways that are consistently up or downregulated in clinical samples relative to 2D culture or 3D microtumours. Enrichment plots show: immunoregulatory interactions that were downregulated in A) 2D cultures and B) 3D microtumours compared to clinical samples; ECM-related genes that were downregulated in C) 2D cultures and D) 3D microtumours compared to clinical samples. Bubble plots show selected pathways that are E) downregulated (-ve NES score) or F) upregulated (+ve NES score) in 2D or 3D microtumours relative to clinical samples.

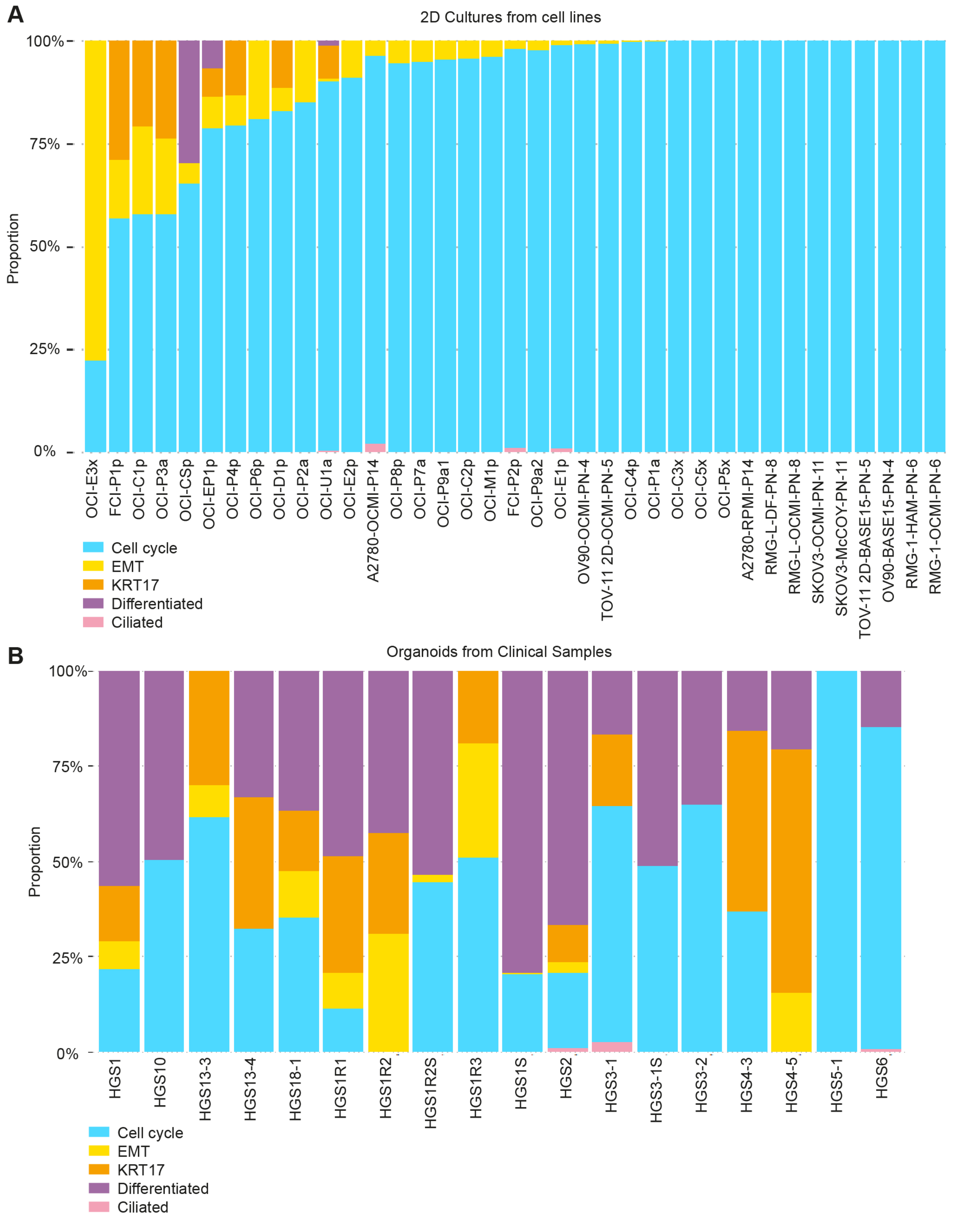


**Figure S7.** Stacked bar plots visualizing the deconvolution result of A) a publicly available dataset of 37 additional ovarian cancer cell lines grown in 2D cultures; B) a publicly available dataset of organoids established from clinical samples. Colours of the bars denote the 5 cell states as shown in the legend.

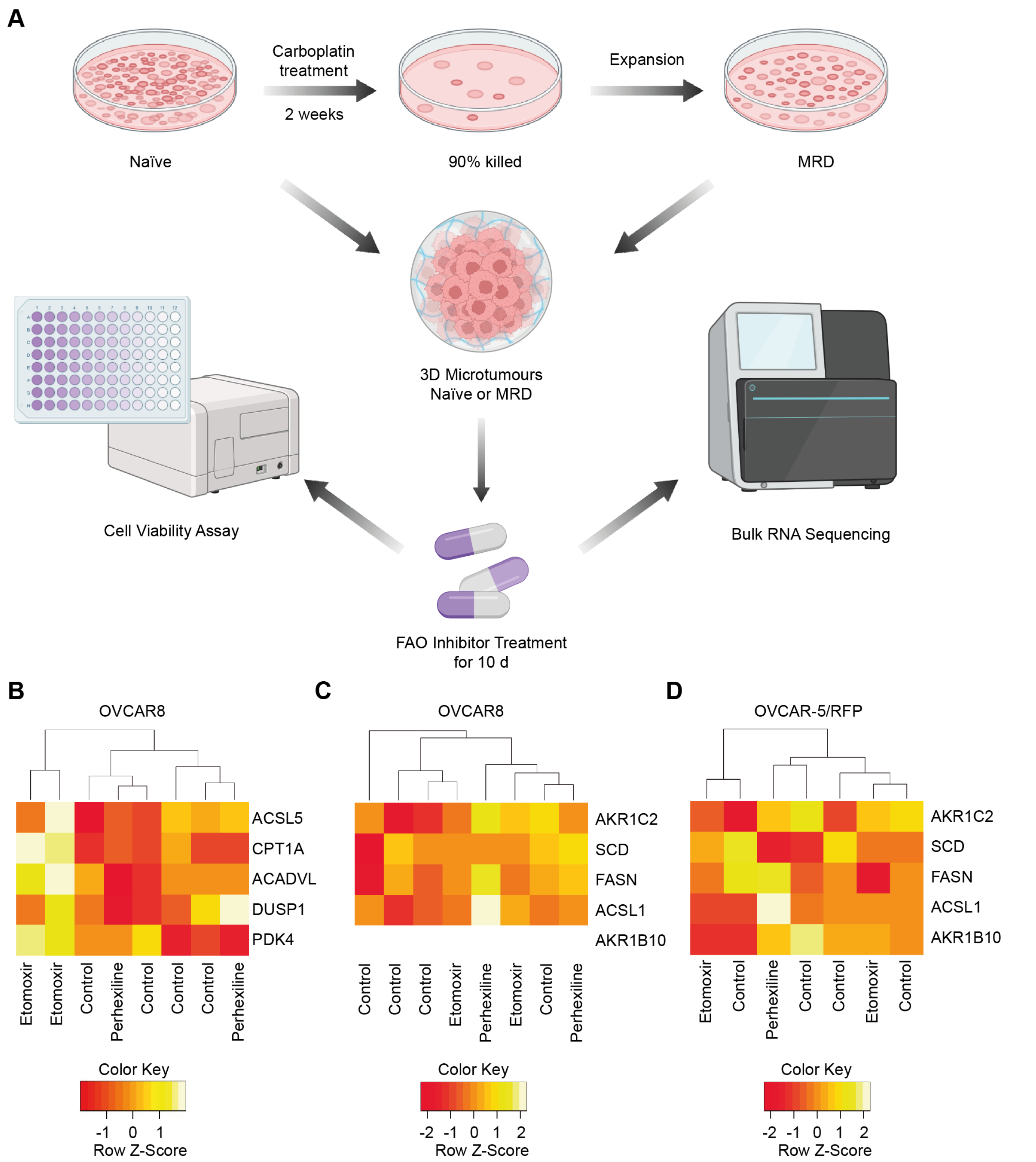


**Figure S8.** A) Schematic diagram of the treatment of naïve and MRD cells with chemotherapy agents. The naïve ovarian cancer cells were treated with carboplatin for 2 weeks to achieve 90% cell killing and the surviving MRD cells were collected and expanded. 3D microtumours were fabricated (D0) from naïve cells or the corresponding MRD cells for each cell line. Next, FAO inhibitor treatment was applied to both types of microtumours for 10 days (D0 to D10). Cell viability assays were performed and mRNAs were extracted for bulk RNA sequencing on D10. The diagrams were created with BioRender.com. B) Heatmap showing differentially expressed genes (DEGs) involved in FAO for etomoxir-resistant MRD 3D microtumours composed of OVCAR8. Heatmap showing DEGs for perhexiline-resistant MRD 3D microtumours composed of C) OVCAR8 and D) OVCAR-5/RFP at D10 (OVCAR8 cells do not express AKR1B10).

**Table S8.** Killing effect comparison between naïve and MRD 3D microtumours.

| Difference in cell viability (compared to DMSO) | | Naïve | MRD | ∆ = Naïve−MRD |
|---|---|---|---|---|
| Etomoxir | OVCAR5 | 30% | -7% | 37% |
| | OVCAR8 | 10% | 12% | -2% |
| | OVCAR-5/RFP | -1% | -6% | 5% |
| Perhexiline | OVCAR5 | -6% | -54% | 48% |
| | OVCAR8 | -6% | -79% | 73% |
| | OVCAR-5/RFP | -15% | -97% | 82% |

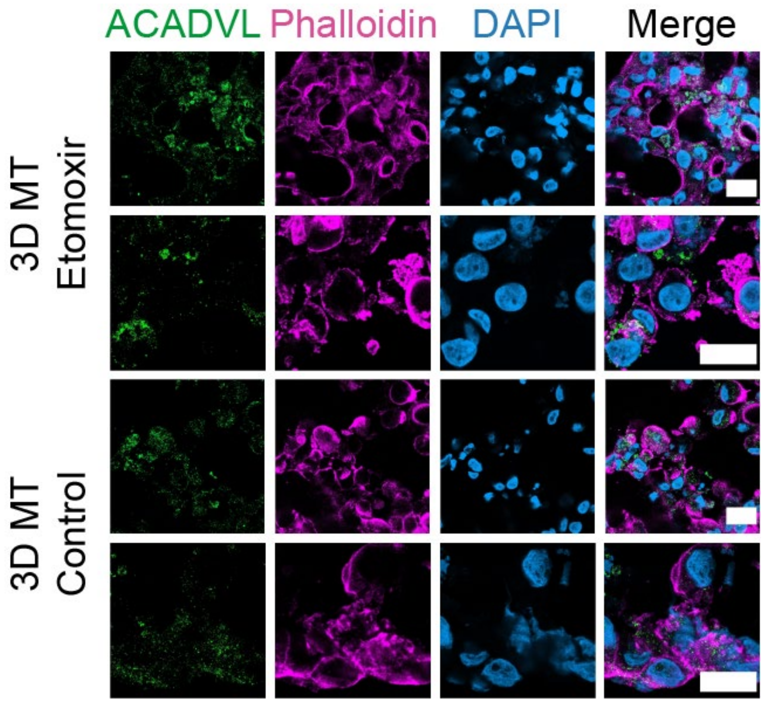


**Figure S9.** Confocal images of MRD 3D microtumours of FAO markers in 3D microtumours at D10: ACADVL (green), phalloidin (pink), DAPI (blue). Scale bar = 20 μm.

**Table S9.** Protocol for differentiation of 3T3-L1 cells into adipocytes.

| Day | Medium |
|---|---|
| D0 | DMEM supplemented with 10% (v/v) bovine calf serum (ATCC, #30-2020) and 1% (v/v) Pen-Strep |
| D2 | DMEM supplemented with 10% (v/v) FBS, 517 µM IBMX (Sigma-Aldrich, #I7018), 1 µM dexamethasone (Sigma-Aldrich, #D2915), and 167 nM insulin (Merck, #407709) |
| D4 | DMEM supplemented with 10% (v/v) FBS and 167 nM insulin |
| D6 | DMEM supplemented with 10% (v/v) FBS |

* Cells were maintained at complete confluence for 2 d (D0 to D2).

**Table S10.** Microplate reader settings for fluorescence intensity measurement to evaluate cell viability.

| Parameter | Value |
|---|---|
| Excitation wavelength | 555 nm |
| Excitation bandwidth | 30 nm |
| Emission wavelength | 595 nm |
| Emission bandwidth | 30 nm |
| Orbital averaging* diameter | 4 mm |

* With the orbital averaging on, measurements were taken within a circle of 4 mm to give the average fluorescence signal of the area. This setting is favoured in case the 3D microtumour does not sit in the centre of each well.

**Table S11.** List of antibodies used for immunofluorescence staining.

| Antibody | Catalogue number | Dilution |
|---|---|---|
| VEGFA | Thermo Fisher MA1-16629 | 1 in 200 |
| NDRG1 | abcam ab124689 | 1 in 200 |
| P4HA1 | abcam ab244302 | 1 in 200 |
| ALDH3A1 | abcam ab76976 | 1 in 100 |
| SCD | Thermo Fisher MA5-27542 | 1 in 100 |
| AKR1B10 | abcam ab96417 | 1 in 500 |
| CPT1A | abcam ab128568 | 1 in 50 |
| ACADVL | abcam ab155138 | 1 in 200 |
| AKR1C1/2 | abcam ab131375 | 1 in 200 |
| FASN | abcam ab128856 | 1 in 50 |
